# The nuclear import receptor Kapβ2 modifies neurotoxicity mediated by poly(GR) in C9orf72-linked ALS/FTD

**DOI:** 10.1101/2022.09.30.510384

**Authors:** ME Cicardi, V Kankate, S Sriramoji, K Krishnamurthy, SS Markandaiah, BM Verdone, A Girdhar, A Nelson, LB Rivas, A Boehringer, AR Haeusler, P Pasinelli, L Guo, D. Trotti

**Affiliations:** Weinberg ALS Center, Vickie and Jack Farber Institute for Neuroscience, Department of Neuroscience, Thomas Jefferson University, Philadelphia, PA, U.S.A; Department of Biochemistry and Molecular Biology, Thomas Jefferson University, Philadelphia, PA, U.S.A

**Keywords:** C9orf72, ALS, FTD, karyopherin β2, dipeptide repeat proteins, TNPO1, TDP-43

## Abstract

Expanded intronic G_4_C_2_ repeats in the *C9orf72* gene cause several cases of amyotrophic lateral sclerosis (ALS) and frontotemporal dementia (FTD). These repeats are translated through a non-AUG-dependent mechanism into five different dipeptides (DPRs), including poly-glycine-arginine (GR), which is aggregation-prone and eventually neurotoxic. Here, we report that Kapβ2 and GR interact, co-aggregating in primary neurons *in-vitro* and CNS tissue *in-vivo*. Importantly, this interaction improves the overall survival of neurons expressing GR. Downregulation of Kap β2 is detrimental to the survival of neurons only if GR is expressed, whereas increased Kap β2 levels mitigate GR-mediated neurotoxicity. notably, we did not find any changes in TDP-43 localization nor in the dynamic properties of the GR aggregates when Kapβ2 was over-expressed. These findings support the design of therapeutic strategies aimed at modulating Kap β2 levels as a potential new avenue for contrasting neurodegeneration in C9orf72-ALS/FTD.

## INTRODUCTION

The most common genetic cause of amyotrophic lateral sclerosis (ALS) and fronto temporal dementia (FTD) is a non-coding G _4_C_2_ nucleotide repeat expansion (NRE) in the first intron of the *C9orf72* gene (Dejesus-hernandez et al., 2011, Renton et al., 2011, Feldman et al., 2022). This NRE can be aberrantly translated through a process known as repeat-associated non-AUG translation (RAN-T), leading to the production of five different dipeptide repeat proteins (DPRs): poly glycine-alanine (GA), poly glycine-proline (GP), poly glycine-arginine (GR), poly proline-alanine (PA), and poly proline-arginine (PR) (Gendron and Petrucelli, 2017). The arginine-rich DPRs, GR and PR, are the most neurotoxic, possibly in a length-dependent manner (Wen et al., 2014, Verdone et al., 2022). Studies in *in-vitro* models have identified dysregulated pathways associated to the accumulation of GR and PR, such as translation stalling, stress granules persistence, nucleolar stress, DNA damage, and nuclear pore complex defects (Coyne et al., 2020, Moens et al., 2019, Maor-Nof et al., 2021, Chew et al., 2019, Boeynaems et al., 2017). A well-known pathological hallmark of most forms of ALS/FTD, including C9-ALS/FTD, (Cook et al., 2020) is the mislocalization of TDP-43 from the nucleus, where it usually resides, to the cytoplasm. Expression of long GR dipeptides (≥200) in cells causes TDP-43 nuclear depletion and accumulation of cytoplasmic TDP43 inclusions. GR_80_-expressing mice show behavioral impairments but lack TDP-43 mislocalization in neurons (Choi et al., 2019). Also, our recently established mouse model, which constitutively expresses GR_50_, develops mild motor behavioral phenotype (Verdone et al., 2022) but fails to display TDP-43 mislocalization in neurons. Interestingly, it displayed disrupt nuclear pore complex integrity, as revealed in aged mice by Nup62 (nuclear pore complex protein) mislocalization in the cytoplasm (Gleixner et al., 2022). From a mechanistic perspective, the influence of GR on nuclear import/export machinery remains unclear. Vanneste et al. showed that *in-vitro* expression of DPRs did not affect nuclear import/export processes (Vanneste et al., 2019), while Hayes et al. showed that R-rich DPRs disrupted the karyopherin-dependent nuclear import in a dose-dependent fashion (Hayes et al., 2020). Hutten et al., addressing specifically TDP-43 shuttling, showed that GR, even at short length (GR_20_), hinders TDP-43 nuclear import (Hutten et al., 2020). In line with this evidence, it was demonstrated that also GA can cause specifically TDP-43 nuclear import defects (Khosravi et al., 2017). TDP-43 mislocalization is strictly a neuronal event, at least in the human disease context. However, these last two studies were conducted in human cancer cell lines (HeLa), whose protein and RNA composition greatly varies from neuronal cells.

The GR interactome is composed of many proteins, most of which feature low complexity domains, through which the interaction with GR occurs, including RNA-binding proteins, nucleolar components, stress granules components, and proteins involved in nuclear transport (Lee et al., 2016, Kwon et al., 2015). Among the latter is karyopherin β2 (Kapβ2, also known as transportin-1 or Importin β2), which belongs to the family of nuclear import receptors (NIRs) that orchestrate the exchange of proteins between the nucleus and cytoplasm (Stewart, 2007). The flow direction depends upon RanGTP/RanGDP asymmetric distribution between the nucleus and cytoplasm. After recognizing and binding to its cargo in the cytoplasm, Kapβ2 translocates to the nuclear envelope (Izaurralde et al., 1997, Dasso, 2002), where it can gain entry into the nucleus through interaction with the FG-Nups on the nuclear pore (Rout et al., 2003, Weis, 2003, Hulsmann et al., 2012, Beck and Hurt, 2017). Once in the nucleoplasm, Kap β2 releases its cargo thanks to the binding with the highly concentrated RanGTP that mediates Kap β2 relocation in the cytoplasm. In the cytoplasm, GTP is dephosphorylated in GDP and thus loses the interaction with Kap β2, which is then ready for a new import cycle (Conti and Izaurralde, 2001, Fried and Kutay, 2003, Weis, 2003).

Even though no ALS-causative mutations in the *KAPB2* gene have been identified thus far, the Kap β2 protein interacts with many ALS-causative proteins such as FUS, but also TAF, EWS, hnRNPA1, and hnRNPA2 (Dormann et al., 2012, Takeuchi et al., 2013, Troakes et al., 2013, Guo et al., 2018). Interestingly, the domain of these proteins involved in the interaction with Kapβ2 is a PY-NLS, a nuclear localization signal in which one proline (P) is followed by one tyrosine (Y), in which many of the most common and aggressive ALS-related mutations are found (Twyffels et al., 2014, Lu et al., 2021, Fare et al., 2023). Even at very short lengths (e.g., 10 repeats), GR is predicted to interact with Kap β2 *via* the same domain (Hayes et al., 2020). A recent study shows that PR also interacts with Kap β2 through the sites that recognize the PY-NLS (Jafarinia et al., 2022). To add importance to the interaction of R-rich DPRs and Kap β2, previous studies in yeasts showed that overexpression of KAP104, the yeast homologue of Kapβ2, decreased PR toxicity (Jovičič et al., 2015), while in flies, Kap β2 RNAi worsens PR neurotoxicity (Boeynaems et al., 2016, Lee et al., 2016).

In this work, we reported that GR interacts with Kap β2 in primary neurons and CNS tissues supporting the relevance to disease of this interaction. We also explored the impact of this interaction on neuronal survival. Indeed, reduced levels of Kap β2 had an adverse effect on neuronal survival in the presence of GR. Conversely, increased Kapβ2 significantly mitigated GR toxicity. These results indicate that Kap β2 could be a positive modifier of a pathogenic disease mechanism. Therefore, manipulations of its expression levels in neurons might be harnessed to design future therapies to improve their survival in C9orf72-linked neurodegeneration.

## RESULTS

### Expression of GR does not alter endogenous levels of Kap**β**2 in neurons

The recruitment of essential constituents of the translational machinery by GR can lead to the stalling of translation, affecting the expression levels of certain proteins (Kriachkov et al., 2022). Thus, we first investigated whether expression of GR could potentially affect Kap β2 protein levels in our experimental models. For experiments in cortical neurons we employed a lentivirus expression vector which encods GR_50_ with Flag and GFP tags fused to the dipeptide to help with visualization and identification. The Flag tag is fused to the N-terminus of GR _50_, while the GFP tag to the C-terminus. In addition, the lentivirus expression vector is driven by the neuronal-specific human synapsin promoter (hSyn). This promoter is active in neurons and will only drive gene expression in these cell types. In cortical neurons transduced with GR_50_-GFP, there was no significant change in Kap β2 mRNA or protein levels compared to neurons expressing GFP used as reference (**Fig 1A-C**). Moreover, we assessed Kap β2 mRNA and protein levels in the cerebral cortex and spinal cord homogenates obtained from a transgenic mouse expressing GR _50_-GFP or GFP (Verdone et al., 2022). This analysis revealed no significant differences between the two groups in Kap β2 mRNA and protein levels (**Fig 1D-F**). We also assessed if there were discrepancies in the expression levels of the GR_50_-GFP and GFP transcripts but did not find significant changes between the experimental groups, both in cortical neurons and mouse CNS tissues (**Suppl Fig 1A-B**). While postmortem tissues of C9orf72-ALS/FTD patients contain GR-positive aggregates, whose abundance varies in different brain regions (Schludi et al., 2015), we found that Kapβ2 protein expression levels were not changing, as determined by Western blot analysis of homogenates obtained from the motor cortex of C9orf72-ALS/FTD and non-diseased control cases (**Fig. 1G, H**). We also analyzed the expression of the Kapβ2 transcripts in publicly available datasets, including the one from the Answer ALS consortium (transcriptomic data from iPS-derived cortical neurons (iCN) from ALS patients) and the ALS Cell Atlas (frontal cortex and cerebellum transcriptomic data from ALS patients and controls) and found no differences in Kap β2 RNA transcript levels when comparing control and C9orf72-ALS/FTD groups (**Suppl Fig 1C, D**).

**Figure 1.**
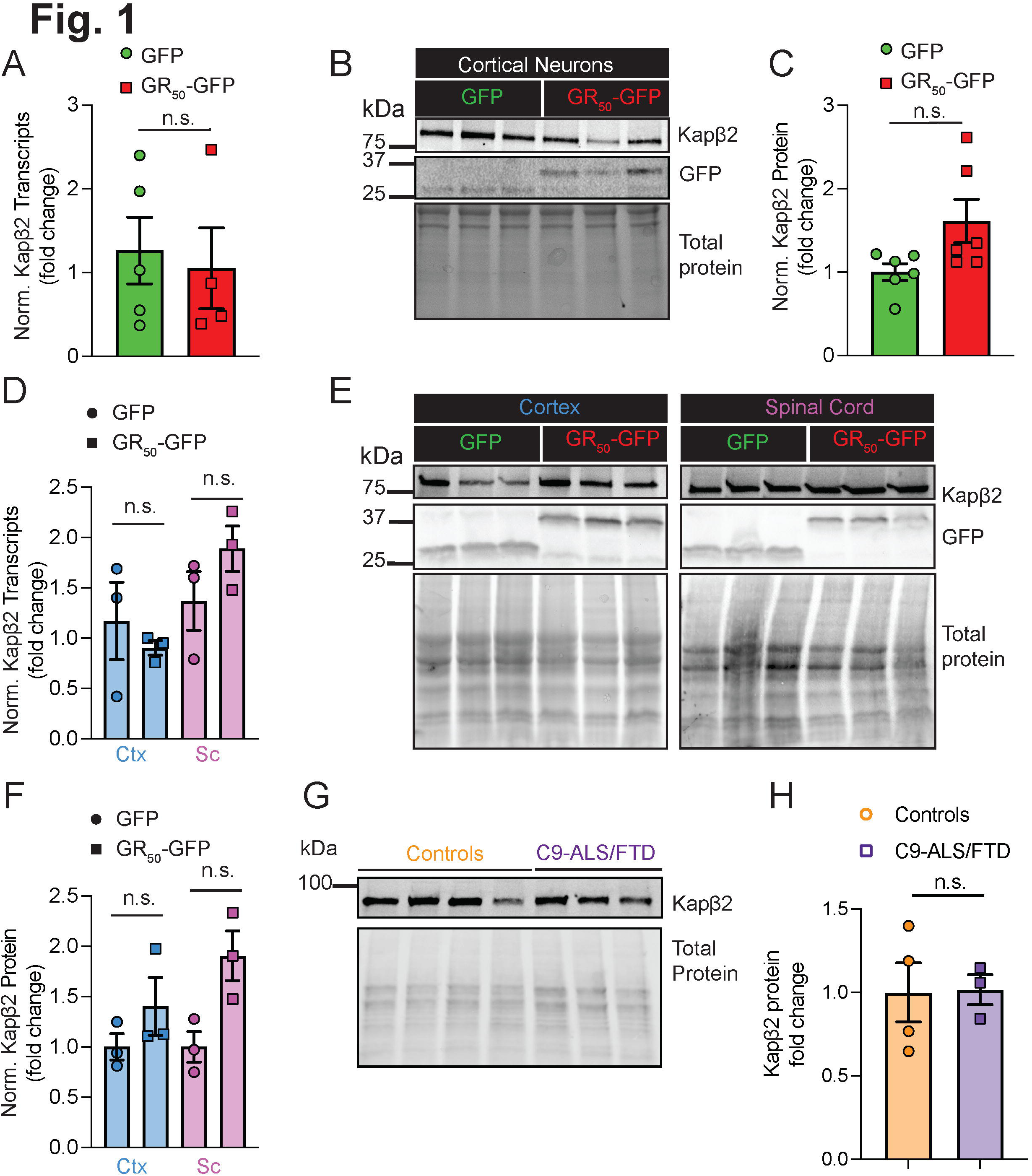
GR does not affect Kapβ2 levels in neurons. A. qPCR analysis of Kapβ2 levels in cortical neurons transduced with GFP or GR_50_-GFP. Data are represented as mean ±S.E.M. (n=6, Student t-test, n.s.) B. Western blot of cortical neuron extracts transduced with GFP or GR_50_-GFP. Blots were probed for GFP and Kapβ2 and total protein was used as loading control. C. Graph bar showing quantification of the WB on cortical neuron extracts for Kap β2 protein. Total protein was used to normalize (n=6, Student t-test, n.s.). D. qPCR analysis of Kapβ2 levels in cortex and spinal cord of GFP and GR _50_-GFP mice. Data are represented as mean ±S.E.M. (n=3, One Way-ANOVA, n.s.) E. Western blot of cortex and spinal cord homogenates from GFP and GR_50_-GFP mice. Blots were probed for GFP and Kapβ2 and total protein was used as loading control. F. Graph bar showing quantification of the WB on cortex and spinal cord homogenates for Kapβ2 protein. Tubulin was used to normalize (n=3, One Way-ANOVA, n.s.) G. Western blot of controls and C9-ALS patient’s motor cortex extracts. Blot was probed for Kapβ2 H. Graphs showing quantification of Kapβ2 expression in controls and C9-ALS motor cortex extracts. Data are represented as mean ±S.E.M. (n=3, Student t-test, n.s.)

### GR interacts with Kap**β**2 in neurons

Earlier investigations utilizing pull-down assays have revealed the interaction between Kap β2 and GR in various human cell line models (Hutten et al., 2020, Hayes et al., 2020). To explore whether this interaction would also occur in cell types relevant to the disease, we employed a lentivirus expression vector to transduce primary cortical neurons with GR_50_. Immunofluorescence staining revealed perinuclear and cytoplasmic localization of Kap β2 in neurons as expected (**Fig. 2A**). To evaluate whether GR _50_ and Kap β2 co-localize and estimate the degree of co-localization of these proteins in the same cellular compartments, we measured the Pearson’s correlation coefficient in transduced neurons. The analysis revealed that the co-localization coefficient was higher than one in GR _50_ neurons, indicating that there is a positive correlation between GR_50_ and Kap β2 (**Fig. 2B**). Next, to investigate whether Kapβ2/GR interacted also in *in-vivo* models, we analyzed a novel GR _50_ transgenic mouse model that we recently generated and characterized (Verdone et al., 2022). This mouse model expresses GR _50_-GFP predominantly in neurons through Cre-recombination. We first assessed the Kapβ2 localization pattern in wild-type, non-transgenic mice and found that Kap β2 mainly localized in perinuclear regions and the cytoplasm of CNS cells (**Suppl. Fig. 2A**). We then performed Kapβ2 immunofluorescence staining in cerebral cortex and spinal cord sections of the GR _50_ transgenic mouse employing Imaris software to perform 3D rendering and visualize the Kapβ2 localization in the cortex and spinal cord neurons. We found robust co-localization of Kapβ2 and GR_50_-GFP (Pearson’s coefficient ranging between 0.5 and 1 in both tissues) in NeuN-positive cells (**Fig. 2C, D; Movie 1**).

**Figure 2.**
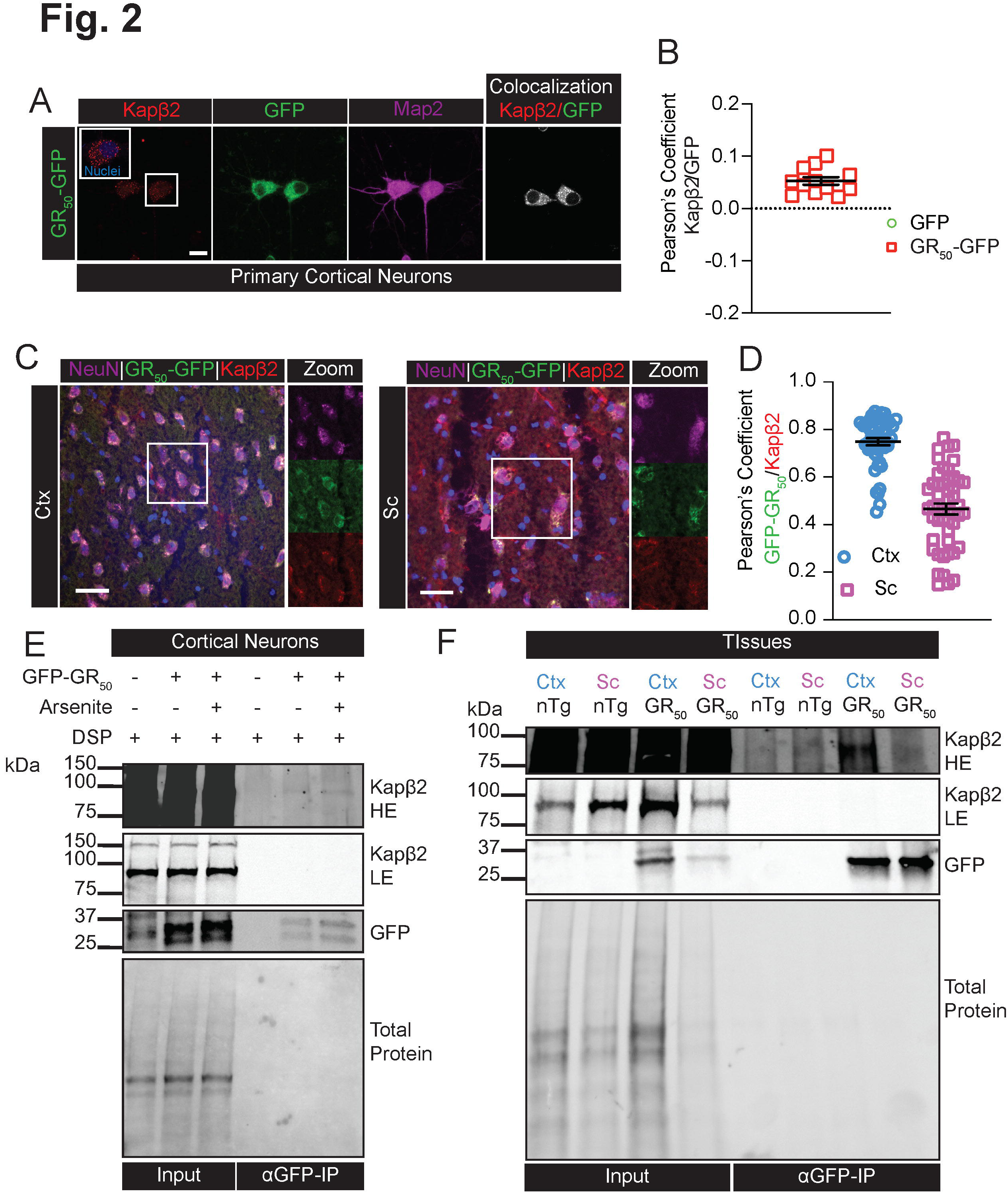
GR interacts and recruits Kapβ2 in neurons. A. Confocal images of primary rat cortical neurons transduced with Flag-GR_50_-GFP (bottom row) and stained for Kap β2. Green is GFP; red is Kap β2, and magenta is Map2. Imaris colocalization shows the amount of GFP colocalized with Kapβ2. Scale bar=10μm. B. Graphs showing Pearson’s coefficient measured in each neuron transduced with GR _50_-GFP. Data are represented as mean ± S.E.M. (n=3, m>5 neurons, Student t-test, P<0.05) C. Confocal images of the motor cortex and spinal cord tissues of GR_50_-GFP mice stained for Kapβ2. Green is GFP; red is Kap β2, and magenta is NeuN. Imaris colocalization shows the amount of GFP colocalized with Kapβ2. Scale bar 30μm. D. Graphs showing Pearson’s coefficient measured in motor cortex and spinal cord neurons of GR_50_-GFP mice. Data are represented as mean ± S.E.M. (n=3, m>15 neurons) E. Western blot of the immunoprecipitation assay performed in rat cortical neurons transduced with GR_50_-GFP and treated with arsenite. Blots were probed for GFP and Kapβ2. F. Western blot of the immunoprecipitation assay performed in the motor cortex or spinal cord tissue of nTg or GR_50_-GFP mice. Blots were probed for GFP and Kapβ2.

Further confirmation for GR _50_ and Kap β2 reciprocal interaction in neurons and CNS tissues could come from co-immunoprecipitation (co-IP) experiments. In our experimental conditions, however, this assay poses a challenge, as the fluorescent reporter GFP could potentially interact with numerous proteins in cells. A dataset of interacting proteins, containing over 2900 entries that were found interacting with both GFP and GR_50_, has been deposited and published (MassIVE identifier MSV000088581) (**Suppl. Fig. 3A**). Since so many proteins were found to interact with both GFP and GR_50_, this raised concerns about the reliability of the co-IP technique for this study. We plotted the intensities of the interacting proteins and found that Kapβ2 (as well as Kapβ2b) were significantly enriched in the GR_50_ group compared to the GFP group (**Suppl. Fig. 3B**). This suggested that Kapβ2 and GR could potential be true interacting partners, despite the GFP tag fused with GR_50_ could potentially have affected the results. The fact that Kap β2 and GR were found to be enriched in the GR _50_ group provides, however, some evidence that they may be *bona fide* interacting partners, although further validation is required to confirm it.

We, thus, carried out co-IP in GR _50_-GFP transduced cortical neurons. Before harvesting the neurons, we treated the sample with the crosslinker DSP to aid preserving even the most dynamic and transient interactions between these two proteins. In immunoprecipitates from GR_50_-GFP neurons, we detected Kap β2, whose interaction with GR_50_ increased in neurons treated with the cellular stressor arsenite (**Fig. 2E**). Co-IP performed with flag-beads (the biochemical tag at the N terminus of GR _50_) also showed the interaction between Kapβ2 and GR (**Suppl. Fig. 3C**). We elected not to use GFP-expressing neurons as control group because of the demonstrated spurious interactions of GFP with multiple proteins, including potentially Kap β2 (**Suppl. Fig. 3A**). Instead, we performed a reverse co-IP experiment using a Kap β2 antibody to capture GR. Our results showed that a fraction of GR immunoprecipitated with Kap β2 in arsenite-treated neurons (**Suppl. Fig. 3D**), providing additional evidence for the interaction between GR and Kapβ2 interaction in neurons.

Co-IP experiments were also conducted on cortex and spinal cord lysates obtained from 12-month-old mice expressing Flag-GR_50_-GFP (Verdone et al., 2022). As seen in neurons *in-vitro*, Kapβ2 co-immunoprecipitated with GR _50_-GFP when we used an anti-GFP antibody but not in non-transgenic mouse CNS tissue or when we used the corresponding control Igg (**Fig. 2F and Suppl. Fig. 3F**). In addition, we performed a reverse co-IP using a Kap β2 antibody, which also revealed evidence of GR bound to Kapβ2 (**Suppl. Fig. 3E**).

### Kap**β**2 is protective against GR-mediated neurotoxicity

After analyzing the interaction between Kapβ2 and GR, we investigated whether this interaction could impact GR toxicity on neurons. We first sought to reduce the expression levels of Kapβ2 in neurons expressing GR and to evaluate the resultant effect on their overall survival. We used a combination of 4 siRNAs specifically designed against the Kapβ2 RNA transcripts. The transfection of these siRNAs resulted in a substantial decrease of Kap β2 protein levels by approximately 50% after 48 hours (**Fig. 3A, B**). Kap β2 regulates the nuclear-cytoplasmic transport of different proteins, including FUS, a well-studied substrate for Kap β2 and an ALS causative protein when mutated (Guo et al., 2018). Surprisingly, knocking down Kap β2 in neurons did not affect nuclear FUS levels, measured by quantifying FUS immunofluorescence signal (**Suppl. Fig. 4A, B**). However, using live-cell imaging microscopy, we estimated a significant reduction in viability in the GR_50_ and GR_100_ (GFP tagged) expressing neurons in which Kapβ2 was silenced (**Fig. 3C**, **Table 1**). We measured the fluorescence intensity of GFP in all experimental groups 48 hours after transfection to determine if Kapβ2 knockdown had any impact on the expression levels of GR _50_ and GR _100_. However, no difference was observed (**Fig. 3D**) even when we looked at nuclear accumulation of GR _50_ and GR _100_ (**Fig. 3E**), suggesting that Kapβ2 is not involved in GR degradation or subcellular localization.

**Figure 3.**
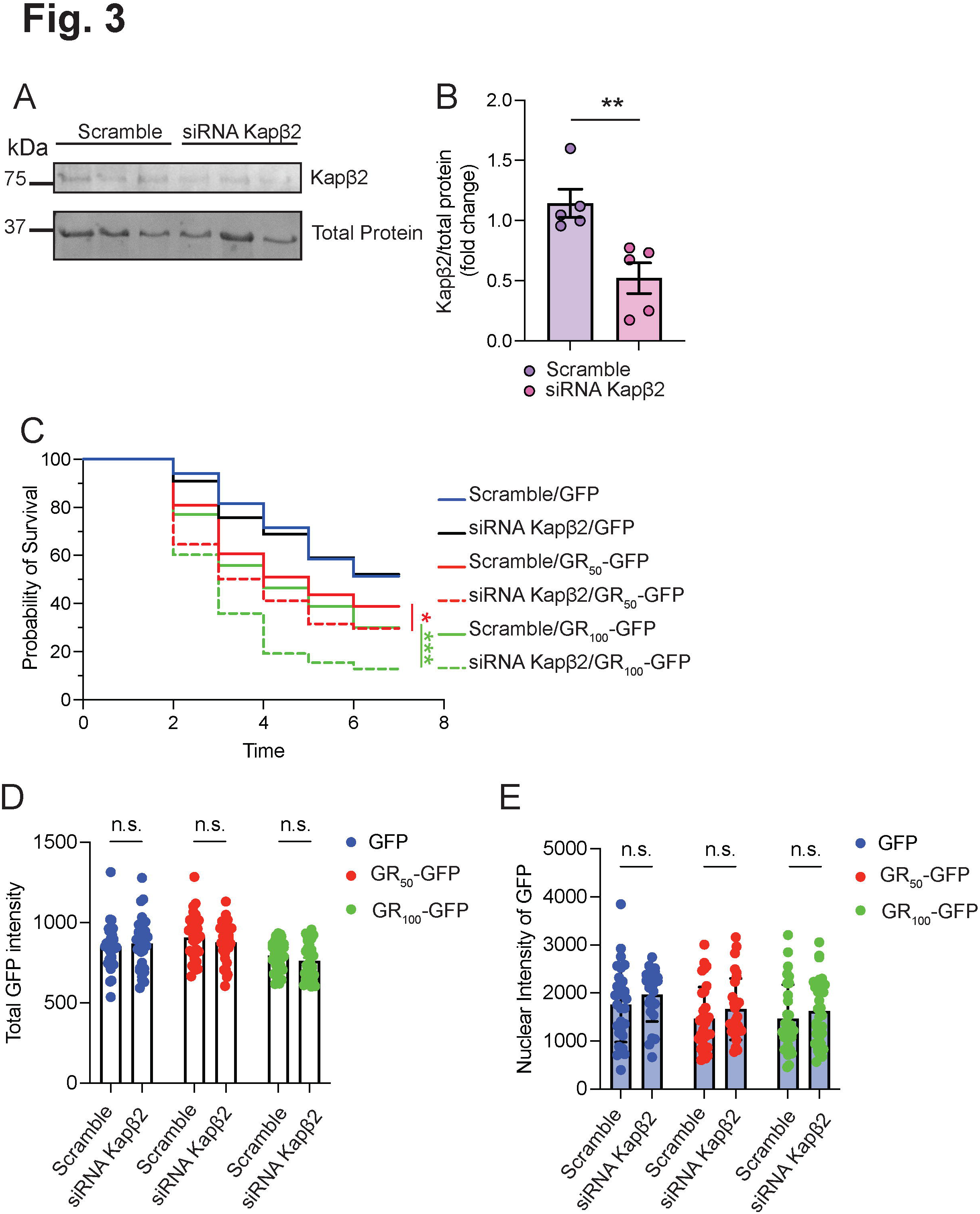
Silencing of Kapβ2 increases death risk in the presence of GR. A. Western blot showing Kapβ2 expression in neurons treated with Smart pool siRNA 100 μM or scramble. B. Graph showing Kapβ2 mRNA in primary neurons treated with Smart pool siRNA 100 μM or scramble. Data are represented as mean ± S.E.M. (n=3, Student t-test, P<0.05) C. Kaplan-Meier curves of time-lapse experiments on rat primary cortical neurons transfected with siRNA Kap β2 100 μM or scramble and GFP, GR _50_-GFP or GR _100_-GFP. (N=3, m>50, Log-rank Mantel-Cox test * p<0.05, *** p< 0.001) D. Quantification of the mean fluorescence of GFP expressing cells 48 hours after transfection in cells previously treated with Kap β2 siRNA or scramble. Data are represented as mean ±S.E.M. (n=3, m>10, One-Way ANOVA, p=n.s.) E. Quantification of the mean nuclear fluorescence of GFP expressing cells 48 hours after transfection in cells previously treated with Kapβ2 siRNA or scramble. Data are represented as mean ±S.E.M. (n=3, m>10, One-Way ANOVA, p=n.s.)

Table 1.

|  | # cells | p-value | Hazard Ratio | 95% CI |
| --- | --- | --- | --- | --- |
| GFP | 222 | 0.0005 | 1.667 | 1.248 to 2.227 |
| GR50-GFP | 188 | Reference Group |  |  |
| GR50-GFP siRNA Kapβ2 | 209 | 0.0137 | 0.7068 | 0.5363 to 0.9314 |
| GFP | 222 | <0.0001 | 2.166 | 1.614 to 2.906 |
| GR100-GFP | 170 | Reference Group |  |  |
| GR100-GFP siRNA Kapβ2 | 98 | 0.0001 | 0.4659 | 0.3167 to 0.6854 |

These results led us to hypothesize that increasing the levels of Kap β2 might play a protective role against GR-mediated neurotoxicity. In order to test the hypothesis, we co-expressed GFP-tagged Kapβ2 and GR _50-100_-mCherry in primary cortical neurons to assess their overall survival. Heterologously-expressed Kap β2 localized in cytoplasmic puncta that mainly surround the nucleus of neurons (**Fig. 4A**, **Movie 2**). We first demonstrated that Kapβ2 or GFP alone were not toxic to neurons when heterologously expressed at two different DNA concentrations (400 and 800 ng/well) (**Fig. 4B, C; Suppl. Fig. 5A, B; Table 2**). This suggested that elevating Kap β2 expression does not have adverse effects on neuronal viability. Next, we expressed both Kap β2 and GR_50_ or GR _100_ and assessed the overall risk of death of the transfected neurons. We kept a 1:1 DNA ratio of Kapβ2-GFP (or GFP alone as control) and GR-mCherry (or mCherry alone), and GFP/mCherry-positive neurons were imaged and analyzed over 8 days (**Fig. 4D, E; Table 3**). As previously reported, expression of GR_50_ or GR_100_ decreased neuronal viability compared to their respective controls (mCherry vs. GR_50_-mCherry p< 0.0001; mCherry vs. GR_100_-mCherry p< 0.0001) as previously reported (Wen et al., 2014b, Verdone et al., 2022). However, co-expression of Kapβ2 in GR _50-100_ (mCherry tagged) neurons decreased their overall risk of death to control levels (GR _50_-mCherry vs. GR _50_-mCherry/Kapβ2 p<0.0001; GR _100_-mCherry vs. GR _100_-mCherry/Kapβ2 p=0.005) (**Fig. 4D, E; Table 3**) indicating that Kapβ2 can indeed rescue neurons from the time-dependent toxicity of GR.

Table 2.

|  | # cells | p-value | Hazard ratio | 95% CI |
| --- | --- | --- | --- | --- |
| GFP - 400 ng | 447 | Reference Group |  |  |
| GFP-TNPO1 - 400 ng | 493 | 0.0714 | 1.18 | 0.9857 to 1.413 |
| GFP - 800 ng | 151 | Reference Group |  |  |
| GFP-TNPO1 - 800 ng | 264 | 0.3556 | 1.13 | 0.8390 to 1.476 |

Table 3.

|  | # cells | p-value | Hazard ratio | 95% CI |
| --- | --- | --- | --- | --- |
| mCherry | 811 | <0.0001 | 0.6796 | 0.5882 to 0.7853 |
| mCherry GR50 | 722 | Reference Group |  |  |
| mCherry GR50/Kapβ2 OE | 520 | <0.0001 | 0.6494 | 0.5538 to 0.7615 |
| mCherry | 811 | <0.0001 | 0.726 | 0.6242 to 0.8444 |
| mCherry GR100 | 623 | Reference Group |  |  |
| mCherry GR100/Kapβ2 OE | 404 | 0.0054 | 0.7791 | 0.6536 to 0.9287 |

**Figure 4.**
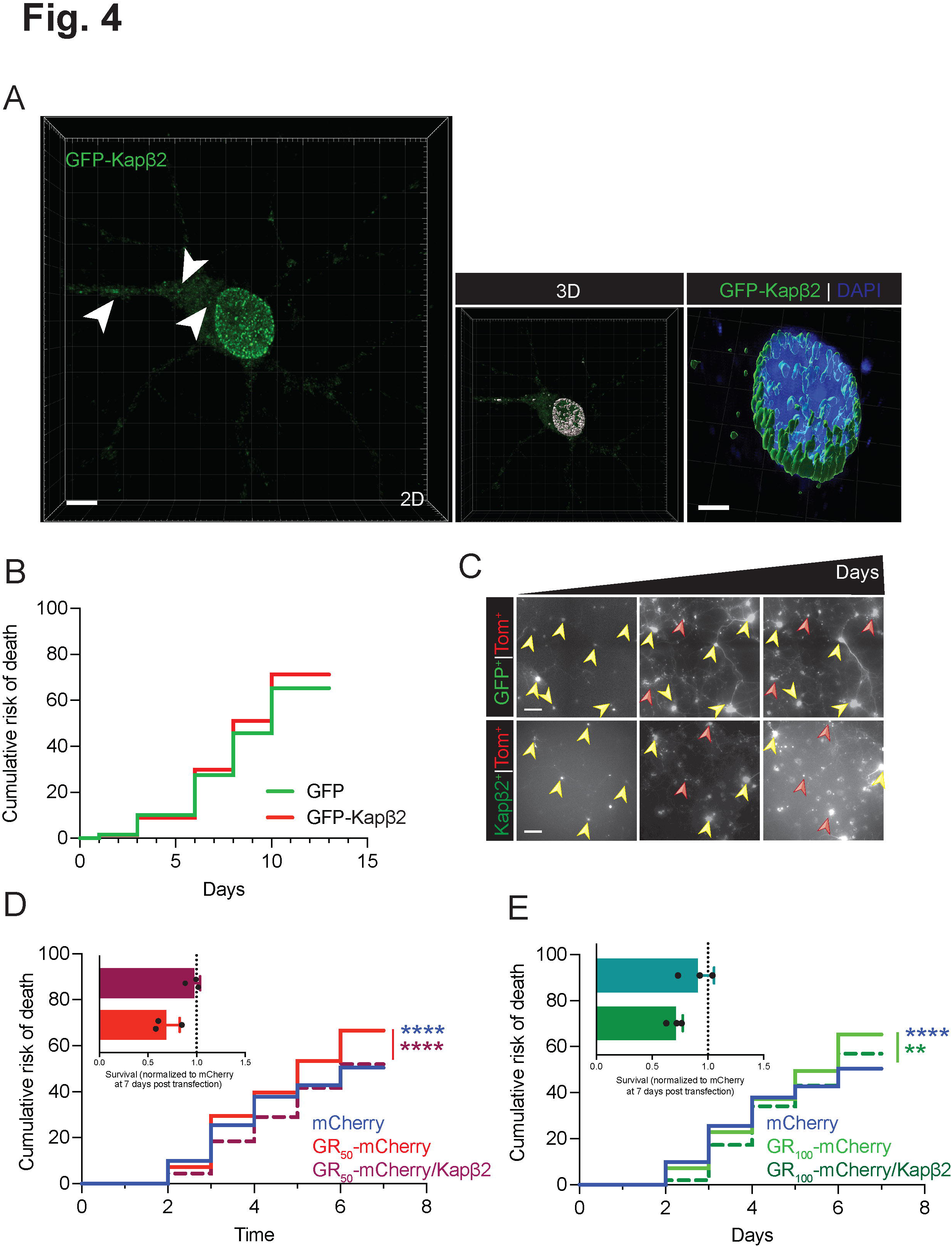
Increased expression of Kapβ2 does not affect neuronal viability. A. Left: Confocal imaging of primary neurons transfected with GFP-Kap β2. Arrows indicate cytoplasmic puncta. Green is GFP, and blue is DAPI. Scale bar 5 μm. Center: Imaris 3D rendering of GFP-Kap β2 expression in primary rat cortical neurons. Right: magnification of Imaris rendering of GFP-Kapβ2 puncta around the nucleus. Scale bar 3μm. B. Kaplan-Meier curves of time-lapse experiments on rat primary cortical neurons transfected with 200/400ng of Tm ^+^/ GFP-Kap β2^+^ (GFP used as control) per 150,000 cells. Neurons double positive Tm ^+^/GFP^+^ were counted (n=3, m>150 neurons, Log-rank Mantel-Cox test: n.s.) C. Representative images of censored neurons over time transfected with 200/400ng of Tm^+^/GFP. Green is Kapβ2; red is Td-Tomato. Scale bar 20μm. D. Kaplan-Meier curves of time-lapse experiments on rat primary cortical neurons transfected with 400/400ng of GFP-Kap β2 (GFP alone used as control) and GR_50_-mCherry (mCherry alone used as control) per 150,000 cells. Neurons double positive GFP ^+^/mCherry^+^ were counted (n=3, m>150 neurons, Log-rank Mantel-Cox test: mCherry vs GR_50_-mCherry p<0.0001; GR_50_-mCherry vs GR_50_-mCherry /GFP-Kapβ2 p<0.0001). Inset: graph bar showing quantification of GR _50_-mCherry and GR _50_-mCherry /GFP-Kap β2 overall survival normalized to mCherry 7 days post-transfection, n=3. Data are represented as mean±S.E.M. E. Kaplan-Meier curves of time-lapse experiments on rat primary cortical neurons transfected with 400/400ng of GFP-Kap β2^+^ (GFP alone used as control) and GR_100_-mCherry (mCherry alone used as control) per 150,000 cells. Neurons double positive GFP ^+^/mCherry^+^ were counted (n=3, m>150 neurons, Log-rank Mantel-Cox test: mCherry vs GR_100_-mCherry p<0.0001; GR _100_-mCherry vs GR _100_-mCherry / Kap β2 p=0.005). Inset: graph bar showing quantification of GR_100_-mCherry and GR_100_-mCherry /GFP-Kapβ2 overall survival normalized to mCherry 7 days post-transfection, n=3. Data are represented as mean±S.E.M.

### Kap**β**2 exhibits length-dependent localization to GR aggregates but doesn’t impact the localization and aggregation of GR in neurons

To dissect the mechanism by which Kap β2 protects in neurons from GR toxicity, we first evaluated whether Kap β2 had an effect on GR aggregates. In neurons, the subcellular localization of GR_50_ and GR_100_ differs significantly. GR_50_ is predominantly located in the nucleus, while GR _100_ is mainly located in the cytoplasm, as shown in **Fig. 5A**. Despite that both GR species form aggregates in the cytoplasm and there was no observed difference in the number of aggregates per cell between GR_50_ and GR_100_ (**Fig. 5E**), we analyzed the intensity profile of the green (GFP reporting on the transfected Kapβ2) and red channels (mCherry reporting on the transfected GRs) in GR aggregates and observed that Kapβ2 exhibited selective localization within the aggregates, as opposed to being diffused in the cytoplasm like GFP, as shown in **Fig. 5A, B**. Furthermore, the fluorescence intensity profile indicated that the recruitment of Kapβ2 to GR aggregates was even higher in GR_100_-expressing neurons, suggesting that Kapβ2 could ultimately affect GR localization. However, we found no difference in GR fluorescence within the nuclei in the presence of Kapβ2, as illustrated in **Fig. 5C**. Additionally, we found that the mean fluorescence of each aggregate remained constant when Kapβ2 was recruited into them, as shown in **Fig. 5D**. Moreover, Kapβ2 did not alter the number of aggregates per cell for either GR species (**Fig. 5E**). Finally, by calculating the Pearson’s correlation coefficient to assess whether the recruitment and colocalization of Kapβ2 and GR depended on the repeat length, we found that GR _100_ aggregates displayed a stronger positive correlation value than GR _50_ (**Fig. 5F, G**).

**Figure 5.**
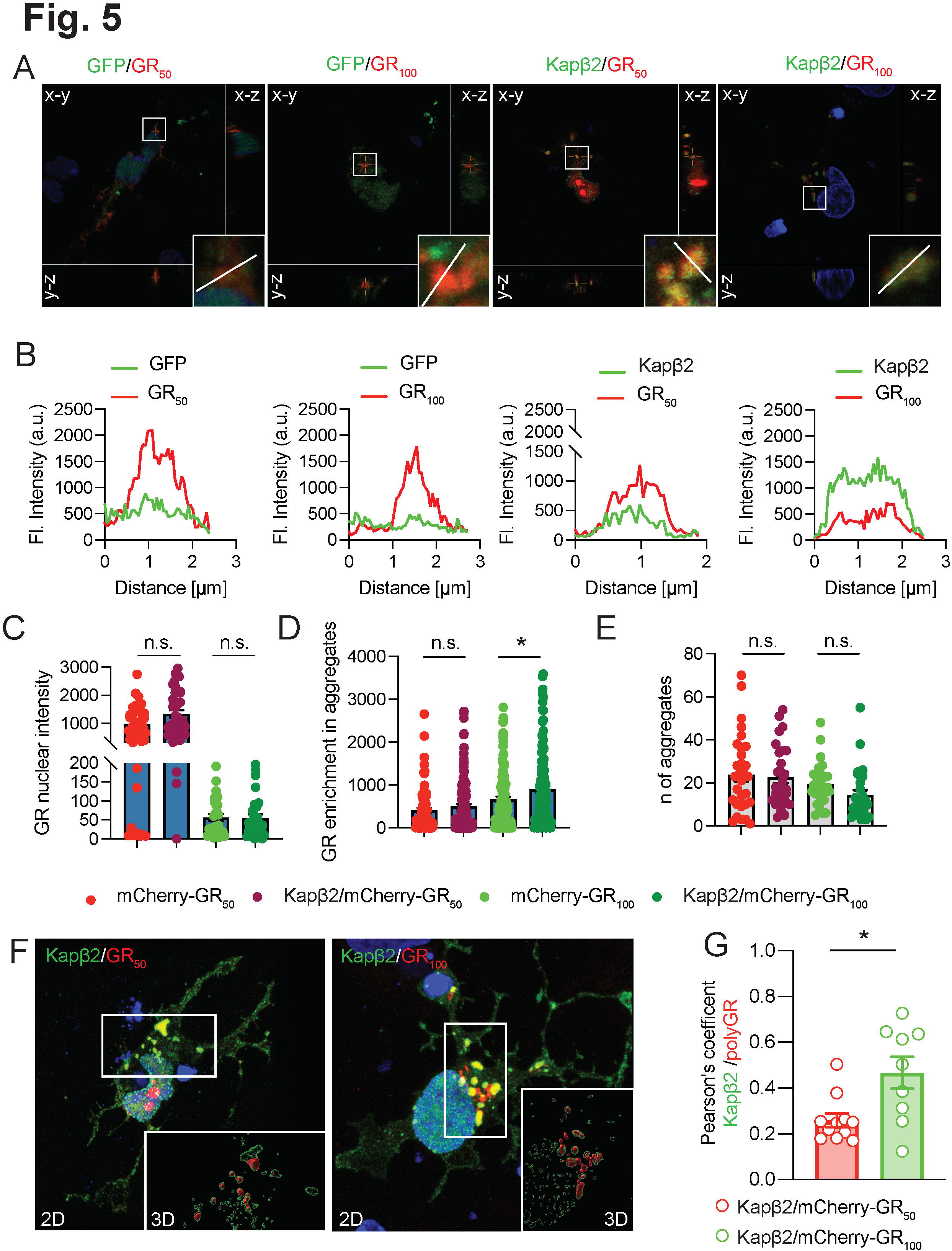
Increased expression of Kapβ2 mitigates GR neuronal toxicity. A. Confocal analysis of neuron overexpressing Kap β2 and GR _50_-mCherry (top row) or GR _100_-mCherry (bottom row). Green is GFP; red is mCherry, and blue is DAPI. The white line represents the length over which GFP and mCherry fluorescence were analyzed. Scale bar 10 μm. B. Intensity profiles are plotted as GFP or mCherry fluorescence (a.u.) over distance (μm). C. Graph bars showing quantification of GR _50_-mCherry and GR _100_-mCherry nuclear intensity Data are represented as mean ± S.E.M. Only double positive cells were considered. (n=2, m>10 neurons, One-Way ANOVA, Tukey’s multiple comparisons test, p=n.s.) D. Graph bars showing quantification of GR_50_-mCherry and GR_100_-mCherry mean fluorescence intensity in GR positive granules. Data are represented as mean ± S.E.M. Only double positive cells were considered. (n=2, m>10 neurons, One-Way ANOVA, Tukey’s multiple comparisons test, p=n.s.) E. Graph bars showing quantification of the number of GR_50_-mCherry and GR _100_-mCherry aggregates per cell. Data are represented as mean ± S.E.M. Only double positive cells were considered. (n=2, m>10 neurons, One-Way ANOVA, Tukey’s multiple comparisons test, p=n.s.) F. Confocal analysis of neuron overexpressing Kap β2 and GR _50_-mCherry (top row) or GR _100_-mCherry (bottom row). Green is GFP; red is mCherry, and blue is DAPI. Imaris rendering is used to visualize the interaction of GFP-Kapβ2 and GRs-mCherry. Scale bar 5 μm. G. Pearson’s coefficient analysis of GFP-Kap β2 and GR _50_-mCherry or GR _100_-mCherry. Data are represented as mean ± S.E.M. (n=2, m>5 neurons, Student t-test, P<0.05)

### Dynamic properties of the GR aggregates are not influenced by Kap**β**2

Kapβ2 affects the dynamic properties of FUS aggregates (Fare et al., 2023), as demonstrated by its ability to modulate their physical properties, liquid phase separation, and formation of insoluble gel-like structures (Guo et al., 2018). To investigate whether Kapβ2 similarly influences the GR aggregates, we co-expressed Kapβ2-GFP and GR_50_ or GR _100_-mCherry in cortical neurons and employed Fluorescence Recovery After Photobleaching (FRAP) to assess the overall fluidity of the aggregates (**Fig. 6A-C**). We observed a higher fraction of mCherry fluorescence recovery for GR_50_-mCherry than for GR_100_-mCherry aggregates, indicating that the aggregates formed by GR _100_-mCherry are less fluid (**Fig. 6A-C**). Notably, in the presence of Kapβ2-GFP, the fraction of mCherry fluorescence recovery remains unchanged for both GR_50_-mCherry and GR_100_-mCherry aggregates, suggesting that Kapβ2 does not bring the aggregates in a more dynamic and fluid state (**Fig. 6D-H**). These findings indicate that Kapβ2 when co-aggregates with GR does not alter the structure of the inclusions aslaready shown in silico by Hutten et al (Hutten et al., 2020).

**Figure 6.**
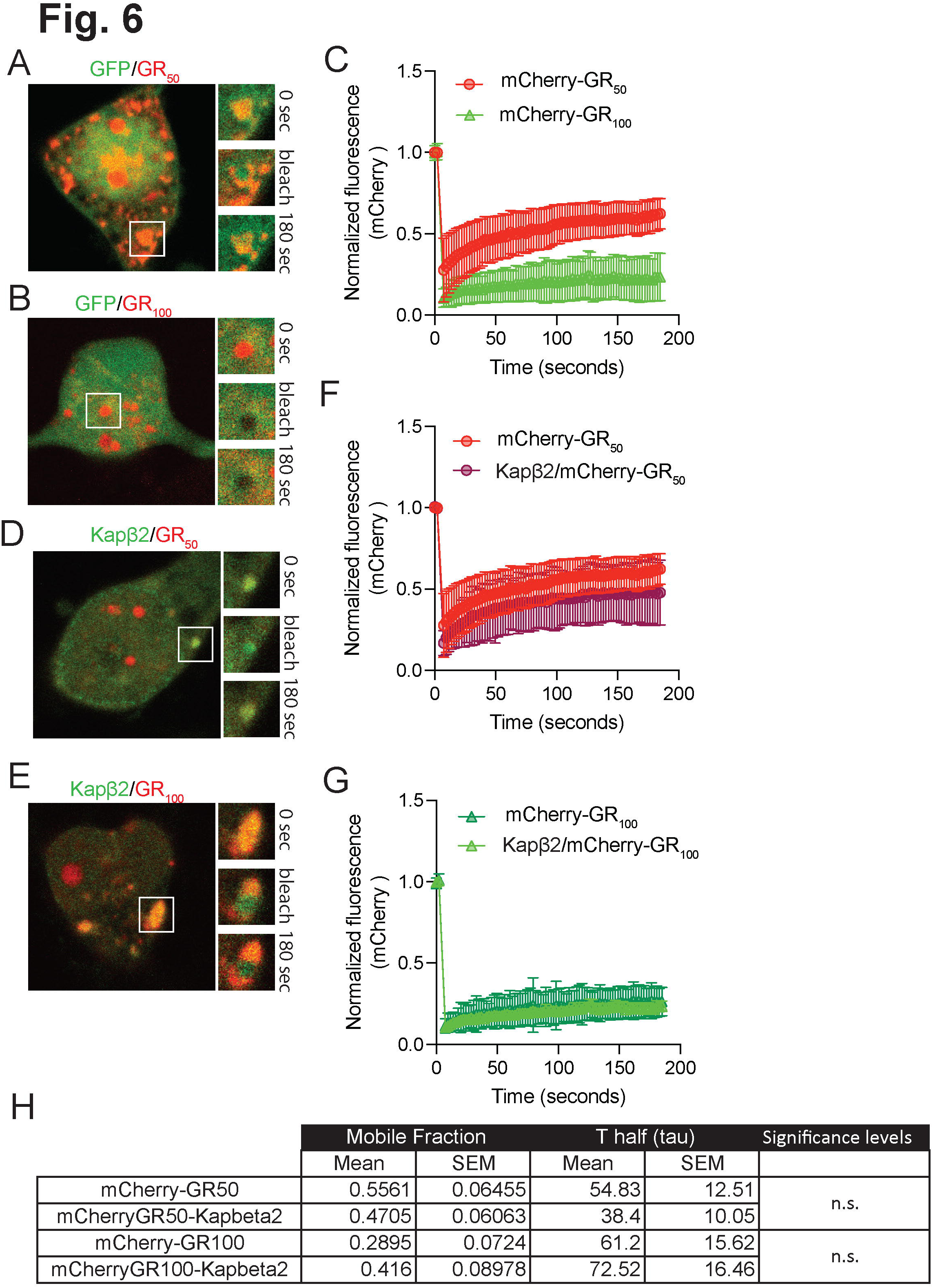
Kapβ2 does not change the dynamic properties of GR aggregates. A. Representative images of the GR_50_-mCherry bleached aggregates at the start (sec=0), bleaching (sec=5), and end (sec=180). Scale bar 10 μm. B. Representative images of the GR_100_-mCherry bleached aggregates at the start (sec=0), bleaching (sec=5), and end (sec=180). Scale bar 10 μm. C. Graph showing fluorescence recovery overtime for GR _50_-mCherry and GR_100_-mCherry. (n=3 m>5, Student t-test, P<0.001) D. Representative images of the GFP-Kap β2/GR_50_-mCherry bleached aggregates at the start (sec=0), bleaching (sec=5), and end (sec=180). Scale bar 10 μm. E. Representative images of the GFP-Kap β2/ GR_100_-mCherry bleached aggregates at the start (sec=0), bleaching (sec=5), and end (sec=180). Scale bar 10 μm. F. Graph showing fluorescence recovery overtime for GR_50_-mCherry and GFP-Kapβ2/GR_50_-mCherry. (n=3 m>5) G. Graph showing fluorescence recovery overtime for GR_100_-mCherry and GFP-Kapβ2/GR_100_-mCherry. (n=3 m>5) H. Table containing mean and S.E.M. for mobile fraction and t half for the represented curves.

### Kap**β**2 does not alter TDP-43 localization in GR-expressing neurons

In presence of GR, Kapβ2 is crucial in regulating TDP-43 localization, although it does not directly bind to TDP-43 (Hutten et al., 2020). We conducted an *in vitro* experiment where GR/TDP-43 aggregates were allowed to form in the presence and absence of Kapβ2. We used an equimolar concentration (5 μM) for all three proteins. When combined *in vitro*, GR and TDP-43 form large condensates (**Fig. 7A, Suppl. Fig. 6**). Interestingly, the addition of Kap β2 reduced aggregates size, TDP-43 enrichment in the aggregates, and increased the diffusive TDP-43 fluorescence signal outside the aggregates (**Fig. 7B, C**), indicating that a fraction of TDP-43 was made soluble by Kapβ2. We then hypothesized that the fraction of TDP-43 freed from the aggregates by Kap β2 would be available to reenter the nucleus freely. We measured intranuclear TDP-43 in neurons co-transfected with GR_50,100_-mCherry and GFP or Kap β2-GFP and found a significant TDP-43 depletion from the nucleus in the presence of GR_50,100_ compared to controls (**Fig. 7D, E**). Under this experimental condition, the expression of Kapβ2-GFP did revert TDP-43 nuclear loss caused by the presence of GR. We thus analyzed TDP-43 content in GR aggregates, and as expected, we found that GR aggregates were enriched in TDP-43.

**Figure 7.**
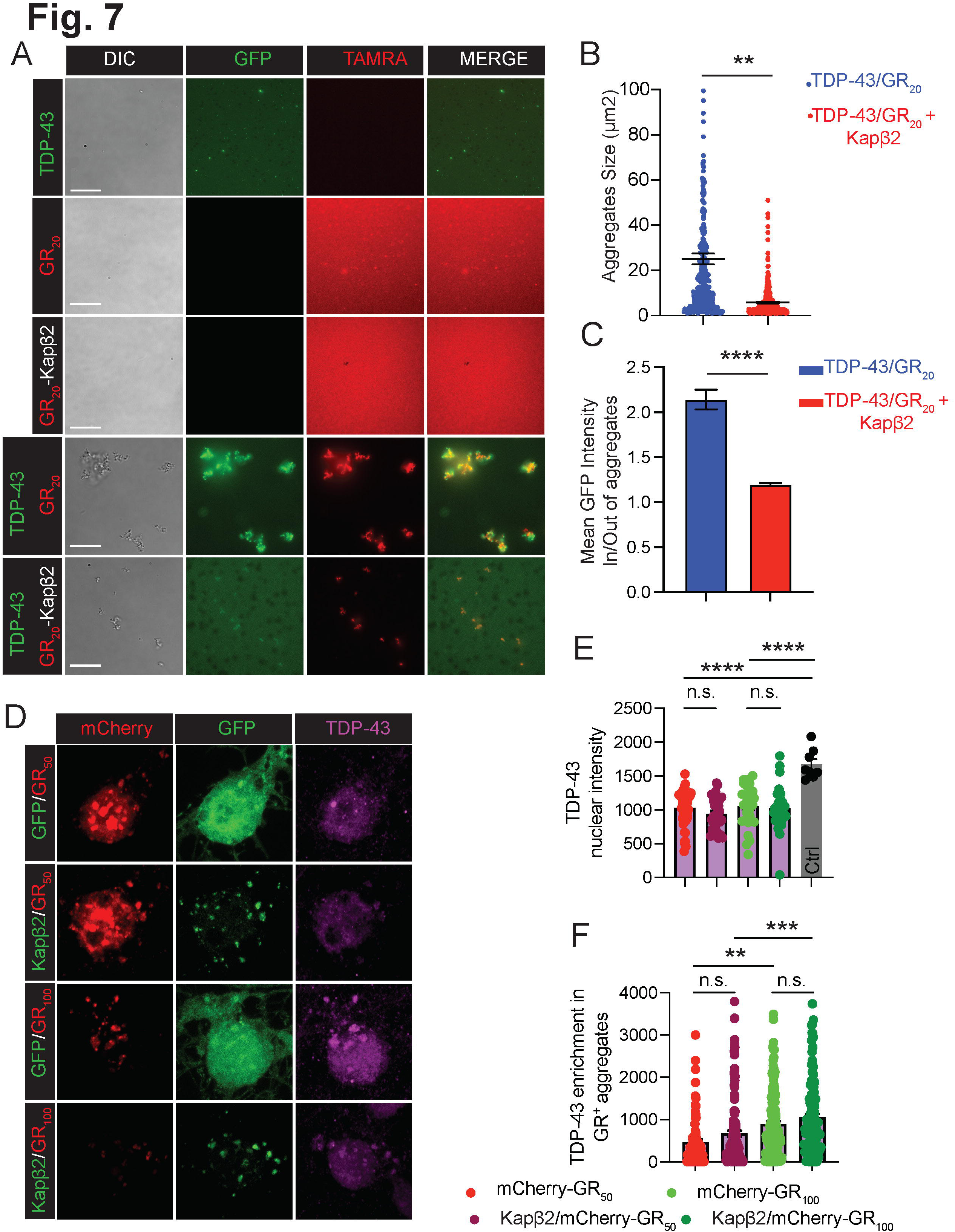
Increasing Kapβ2 expression levels does not affect TDP-43 subcellular localization in the presence of GR. A. Epifluorescence images of condensates formation in the presence of TDP-43, (GR)_20,_ and Kapβ2. Green is TDP-43, and red is GR_20_. Scale bar: 15μm. B. Graph showing quantification of the size of TDP-43/GR20 aggregates in the presence or absence of Kapβ2. Data are represented as mean ± S.E.M. (n=3, Student t-test, P<0.01) C. Graph showing the ratio between the mean GFP intensity inside and outside TDP-43/GR20 aggregates in the presence or absence of Kap β2. Data are represented as mean ± S.E.M. (n=3, Student t-test, P<0.0001) D. Confocal analysis of neuron overexpressing GFP or Kap β2 and GR _50_-mCherry (top row) or GR_100_-mCherry (bottom row). Green is GFP, red is mCherry, magenta is TDP-43, and blue is Hoechst. Scale bar 5 μm. E. Graphs showing single-nuclei TDP-43 staining quantification. Data are represented as mean ± S.E.M. Only double positive cells were considered. (n=2, m>10 neurons, Student t-test, P<0.01; P<0.001) F. Graphs showing TDP-43 quantification into GR aggregates. Data are represented as mean ± S.E.M. Only double positive cells were considered. (n=2, m>10 neurons, Student t-test, P<0.01; P<0.001)

However, we also found that the fraction of TDP-43 in the aggregates did not change in the presence of Kap β2 (**Fig. 7D-F**). Overall, while Kapβ2 makes GR aggregates more dynamic in neurons, it does not influence TDP-43 relocalization to the nucleus or the aggregates.

## DISCUSSION

This study demonstrated the impact of modifying Kapβ2 levels on GR-mediated neuronal toxicity. We established that GR interacts with Kapβ2 *in-vitro* in primary neuronal culture and *in-vivo*, in CNS tissues of a GR-expressing mouse model. Both Kapβ2 and GR have been implicated in the pathogenesis of certain forms of amyotrophic lateral sclerosis (ALS) and frontotemporal dementia (FTD). Kap β2 is a nuclear import receptor that plays a role in regulating the transport of molecule substrates between the nucleus and the cytoplasm.

Dysregulation of nuclear-cytoplasmic transport is thought to contribute to the pathogenesis of ALS and FTD, and it is possible that an interaction between Kap β2 and GR could have implications for this process. Indeed, we found that suppressing Kapβ2 levels elevates the risk of neuronal death. On the other hand, Kapβ2 overexpression shields neurons against GR-induced neurodegeneration.

The formation of GR aggregates is believed to be linked to several cellular dysfunctions in C9orf72-ALS/FTD, including impaired nucleocytoplasmic transport, proteostasis, and RNA metabolism. Dysregulation of these cellular processes can lead to the accumulation of toxic protein inclusions, including GR aggregates, which can trigger a cascade of toxic events that lead to neuronal dysfunction and death. We demonstrated that Kapβ2 exhibited selective localization within GR aggregates but that its presence does not increase the mobile fraction of GR within the aggregates. An increased mobility of the aggregated proteins is normally required for a better resolution of their aggregation state since other proteins, such as proteins involved in inclusion clearance mechanisms, might have access to the aggregates and extract the proteins that could be targeted for degradation. This can facilitate the disassembly and degradation of the aggregates freeing co-aggregated proteins essential for cell survival such as TDP-43 (Lim and Yue, 2015). We were thus not surprised to observe that Kapβ2 did not alter TDP-43 localization in the GR aggregates. TDP-43 mislocalization is a pathological feature of the majority of the ALS cases. It refers to the abnormal accumulation or redistribution of TDP-43 in cytoplasmic inclusions in affected neurons and its depletion from the nucleus, where it physiologically resides (Neumann et al., 2006). This mislocalization triggers a cascade of toxic events that lead to neuronal dysfunction and death (Melamed et al., 2019, Ma et al., 2022).

Hence, potential therapeutic strategies for ALS could be directed to prevent TDP-43 mislocalization and restore its nuclear localization. Several studies, including the present one, have also demonstrated a physical interaction between GR and TDP-43. For example, a study by Chew et al. (Chew et al., 2019) showed that GR directly interacted with TDP-43 and sequestered it in cytoplasmic aggregates, leading to its depletion from the nucleus. This mislocalization of TDP-43 is thought to be a key step in the pathogenesis of ALS and FTD. In our *in-vitro* assay, we found that the addition of Kapβ2 hindered the entry of TDP-43 into the GR aggregates and allowed TDP-43 to remain soluble in the buffer. However, we failed to validate this mechanism in primary neurons in culture. Indeed, we did not observe an increase in nuclear TDP-43 re-localization. Interestingly, these results contradict the previous findings by Hutten et al., which showed that Kapβ2 could directly disrupted GR/TDP-43 co-aggregation, restoring TDP-43 nuclear localization. The ability of DPRs to disrupt the nucleo-cytoplasmic transport and indirectly promote TDP-43 aggregation was also shown by a recent study in HeLa cells (Ryan et al., 2022, Khosravi et al., 2017). In our neuronal culture model, we observed co-aggregation of TDP-43 with GR_50_ and GR_100_ and a corresponding decrease in nuclear TDP-43. While Kapβ2 also localizes to the GR aggregates, it does not reduce TDP-43 localization in the aggregates or alter the TDP-43 nucleo/cytoplasmic ratio.

Numerous studies in the ALS/FTD field have focused on the disruption of the nuclear pore complex and the nuclear import/export activity (Zhang et al., 2015, Coyne et al., 2020, Dubey et al., 2022). Defects in the proper import/export of critical proteins are associated with DPRs or G _4_C_2_ repeat-containing C9orf72 RNA transcripts, while TDP-43 aggregates trigger defects in nucleo-cytoplasmic transport, highlighting the importance of these pathways in disease (Hayes et al., 2020, McLean et al., 2014). Therefore, conducting studies that further characterize these pathways is crucial. A recent study sheds light on the potential consequences of R-rich DPR interaction with the nuclear import receptors (NIRs), proteins responsible for transporting molecules in and out of the cell nucleus through the nuclear pore complex (Nanaura et al., 2021). The study reveals that NIR recruitment into DPR aggregates impairs their ability to maintain the solubility of other RNA-binding proteins (RBPs), such as FUS. Even though it is outside the scope of this study to analyze the behavior of the different RBPs, in the case of TDP-43 we did not observe TDP-43 aggregates other than the ones formed by co-aggregation with GR and Kapβ2.

The pathogenesis of C9orf72-associated ALS/FTD is likely multifactorial, involving multiple mechanisms that interact with each other. A Kap β2-based therapeutic strategy might not bring enough benefits as a stand-alone treatment for ALS but might work in conjunction with drugs targeting other pathways. Kapβ2 levels could conceivably be raised in neurons using a gene therapy approach such as mRNA-nanoparticle technology (Hou et al., 2021). Future studies would also be directed to screening for compounds that can induce the expression of Kapβ2 in neurons. Another intriguing therapeutic possibility is *via* promoting arginine methylation. The Kapβ2-FUS interaction is modulated by FUS methylation status (Hofweber et al., 2018). Arginine methylation is a GR post-translational modification that sometimes dictates disease severity (Gittings et al., 2020). The most abundant GR methylation status in post-mortem tissues is the asymmetric di-methylated form, but GR can be present with symmetrical di-methylation or mono-methylation. It would be essential to expand on this concept and study how the different methylation status/conditions of GR influence Kap β2 binding and, thus, GR toxicity. These studies could entail evaluating the ideal dosage and delivery modes of Kapβ2 and any possible adverse reactions and enduring impacts on the brain.

## ACKNOWLEDGMENTS

We are grateful to the Jefferson Weinberg ALS Center members for their feedback and support throughout the development of this project. The following sources provided funding for this work: National Institutes of Health R21-NS090912 (D.T.), RF1-AG057882 (D.T.), R01-NS109150 (P.P.); Muscular Dystrophy Association grant 628389 (D.T.); DoD grant W81XWH-21-1-0134 (M.E.C.); RF1-NS114128 (A.H.); Family Strong for ALS & Farber Family Foundation (A.H., P.P., D.T.); R35GM138109 (L.G.), Dr. Ralph and Marian Falk Medical Research Trust (L.G.), Frick Foundation for ALS Research (L.G.)

## AUTHOR CONTRIBUTION

M.E.C. and D.T. conceived the study, designed the experiments, interpreted the results, and wrote and revised the manuscript. D.T. supervised all the experiments and provided the resources for the project. M.E.C. performed the live-cell imaging experiments and the statistical analysis. S.S.M. and L.B.R. performed FRAP experiments and analyzed the live-cell imaging data. V.K. performed the experiments in mouse tissues, the co-immunoprecipitations, and the TDP-43 localization experiments. B.M.V. managed the animal colony. S.M. performed the experiments with human samples. K.K. prepared primary neuronal cultures. A.B. performed Imaris analysis. L.G. and A.G. performed the experiments in the test tube and conducted data analysis. A.T.N. designed lentiviral plasmids and produced the virus.

## MATERIALS AND METHODS

### Human tissues

Human post mortem CNS tissue from control individuals and C9orf72 patients were obtained from the Target ALS and the Jefferson Weinberg ALS center biobanks. Information and demographic about the patients are in Table 4.

Table 4.

| Patient | R/G | Yr of Death | age at DOD | age at onset | ATPSY INTERV | Site of Onset | approx. duration in months | Diagnosis | Notes |
| --- | --- | --- | --- | --- | --- | --- | --- | --- | --- |
| 103 | M | 2015 | 22 |  | 9HRS |  |  | Ctrl | Metastatic synovial sarcoma |
| 101 | F | 2015 | 70 |  | 32HRS |  |  | Ctrl | Respiratory failure secondary to lung cancer and infection |
| 95 | M | 2013 | 72 |  | 7HRS |  |  | Ctrl | Cardiorespiratory failure; Pneumonia, myodysplastic syndrome, acute renal failure |
| JWAC 06 | C/F | 2018 | 91 |  | 1 day |  |  | Ctrl | N/A |
| JWAC 29 | C/M | 2019 | 55 |  | 2 days |  |  | C9orf72 |  |
| JWAC 11 | C/M | 2019 | 58 |  | 1 day |  |  | C9orf72 |  |
| JWAC 104 | C/M | 2023 | 53 |  | 1 day | Bulbar |  | C9orf72 |  |

### Cell cultures

HEK293 cells were cultured in DMEM medium (Cytiva Cat.# SH30243.LS) supplemented with 10% FBS (Cytiva Cat.# SH30071.01HI), penicillin, and streptomycin (Thermo Fisher Scientific Cat.# SV30010). Cells were passaged every 3-4 days using 0.05% trypsin (Corning Cat.# 25-051-C).

### Primary cortical neurons

After meninges removal, brains were dissected from embryonic day 16 (E16) rat embryos. Cortices and midbrain regions were cut into small pieces and incubated on a shaker at 80 rcf for 45 min at 37°C in 0.2% trypsin in HBSS without Ca^2+^ and Mg ^2+^ (Cytiva Cat.# SH30588.01). FBS (Cytiva Cat.# SH30071.01HI) was added, and the cell suspension was centrifuged at 800 rcf for 10 min. at 4 °C. Cells were washed with Ca ^2+^ and Mg ^2+^ free HBSS and centrifuged at 800 rcf for 10 min. at 4 °C. The cell suspension was then passed through a 70 μm strainer (Foxx Life Sciences Cat.# 410-0002-OEM) to remove undigested connective tissue and large cell clumps. Dispersed cells were then counted and plated in poly-D lysine-coated plates. The following plating densities were used: 150,000 cells/well in 24 well plates; 300,000 cells/well in 12 well plates; 5,000,000 cells/dish in 10 cm dishes. Arsenite 0.5mM was applied for 1 hour immediately before harvesting. DSP (dithiobis(succinimidyl propionate)) was added to the neurons before harvesting at a concentration of 0.1mM.

### Animals

Flag-GR_50_-GFP (GR_50_-GFP) or Flag-GFP (control-GFP) mice were generated as described. At 12 months of age, mice were sacrificed, and cortices and spinal cords were collected and frozen in liquid N_2_ or embedded in OCT. Frozen samples were stored at –80°C till the time of processing.

### Plasmids and siRNA

The following plasmids were used: pCDNA3_hSyn_Td_Tomato; pCDNA3_hU6_Flag_mCherry; pCDNA3_hU6_Flag_GR_50__mCherry; pCDNA3_hU6_Flag_GR_100__mCherry; pCDNA3_CMV_eGFP; pCDNA3_CMV_eGFP_Kapβ2. eGFP and eGFP_Kapβ2 were co-transfected with Td_Tomato at a DNA concentration ration of 1:2 (200ng/400ng) or 1:4 (200ng/800ng). eGFP and eGFP_Kap β2 were co-transfected with mCherry, mCherry_GR_50_ and mCherry_GR_100_ at 1:1 ratio (400ng/400ng).

Human TDP-43 and TDP-43-GFP were subcloned into pE-SUMO (LifeSensors, Malvern, PA) as described (McGurk et al., 2018). All plasmid inserts were sequenced and confirmed to be correct.

To silence Kap β2, 50 μM of ON-TARGETplus siRNA-TNPO1-Smart Pool (Horizon Discovery, L-086861-02-0005) was used. 50 μM of AllStars Negative Control scramble siRNA (Qiagen Cat.# 1027280) was used as control.

### Transfection

Lipofectamine 2000 (Fisher Scientific Cat.# 11-668-019) was used to transfect HEK cells and cortical neurons (1 μL Lipo2000/500ng total D.N.A.). HiPerfect Transfection Reagent (Qiagen Cat.# 301705) was used to transfect siRNA (8 μL HiPerfect/50nM siRNA). siRNA was transfected 24h before plasmids transfection. For cortical neurons, Lipo/DNA or RNA mix was incubated with the neurons for 1h, and then cell media were changed with fresh ones. For HEK293 cells, the mixture was incubated with the cells overnight.

### Lentivirus constructs production

HEK293 cells were transfected with the following plasmids: pLenti_hSyn_Flag_GR_50__eGFP, psPAX2 (Addgene Plasmid #12260), pMD2.G (Addgene Plasmid #12259). Plasmids were diluted in DMEM, and PEI MAX was added as a transfection reagent.

The mix was incubated with HEK293 cells confluent at 70% for 5h, the media was changed with a fresh one. Cells supernatant was collected 48h after transfection and centrifuged for 10 min at 2,000 rcf at 4°C. Lentivirus Concentrator (OriGene Technologies Cat.# TR30025) was added to the supernatant and incubated overnight at 4 °C. To pellet the virus, the media was centrifuged at 500 rcf for 45 min at 4 °C. Viral particles were then resuspended in Ca ^2+^ and Mg ^2+^ free PBS (Cytiva Cat.# SH30028. L.S.), aliquoted and immediately frozen in liquid N_2_.

### Purification of Recombinant Proteins

His_6_-SUMO1 N-terminally tagged TDP-43-WT and TDP-43-GFP were purified as described (McGurk et al., 2018) and overexpressed in BL21(DE3) RIL *E.coli* cells. These cells were then sonicated on ice in 50 mM HEPES (pH 7.5), 2% TritonX-100, 300 mM NaCl, 30 mM imidazole, 5% glycerol, 2 mM β-mercaptoethanol, and protease inhibitors (cOmplete, EDTA-free, Roche). The recombinant TDP-43 proteins were purified over Ni-NTA agarose beads (Qiagen) and eluted using 50 mM HEPES (pH 7.5), 150 mM NaCl, 300 mM imidazole, 5% glycerol, and 5 mM DTT. The buffer was exchanged with the same buffer without imidazole. The eluate containing the recombinant TDP-43 proteins was then frozen in liquid N _2_, and stored as aliquots at –80°C until use.

Kapβ2 was purified as described (Guo et al., 2018). In brief, *E. coli* BL21-CodonPlus(DE3)-RIL cells (Agilent) were transformed with GST-Tev-Kap β2 plasmid, and expression was induced overnight at 25°C with 1 mM IPTG. Cells were pelleted and resuspended in Tris buffer (50 mM Tris pH 7.5, 100 mM NaCl, 1 mM EDTA, 20% (v/v) glycerol, 2 mM DTT, supplemented with protease inhibitors), then lysed by sonication. Cell lysate was then loaded onto glutathione Sepharose^TM^ 4 Fast Flow resin (GE Healthcare), and washed with Tris buffer, followed by ATP buffer (50 mM Tris pH 7.5, 100 mM NaCl, 1 mM EGTA, 0.5 mM MgCl _2_, 5 mM ATP, 20% glycerol, 2 mM DTT, supplemented with protease inhibitors), then washed and eluted with Buffer A (20 mM imidazole, 75 mM NaCl, 1 mM EDTA, 20% (v/v) glycerol, 2 mM DTT). Finally, the protein was cleaved with Tev protease and purified on a HiTrap Q HP column (GE Healthcare) using a salt gradient. Purified protein was concentrated, flash frozen, and stored at –80 °C. Before the aggregation assay, the protein was thawed and centrifuged at 16,100 x *g* for 10 min to remove any preformed aggregates. Protein concentration was determined by Bradford assay (Bio-Rad, Hercules, CA).

Equimolar concentrations of Kap β2, TDP-43, and (GR) _20_ (5μM) were used. 200nM HIS _6_-SUMO-TDP-43-GFP and 100nM TAMRA-(GR)_20_ were added to visualize TDP-43 and (GR)_20_.

### Peptides

Chemically synthesized (GR) _20_ and Tetramethylrhodamine (TAMRA)-tagged (GR) _20_ were obtained as a lyophilized powder from Peptide2.0 and dissolved in PBS.

### Sample preparation

For human tissues: 100 mg of fresh frozen lumbar spinal cord samples were homogenized in 1% SDS using a 15ml Dounce Homogenizer. The homogenate was centrifuged at 3000 rcf for 20 minutes at 4°C to remove tissue debris. Protein estimation was performed on the clear homogenate using a BCA assay (Pierce BCA kit# 23225). 30 μg of protein samples are loaded for Western blot analysis.

For mouse tissues: 50 mg of CNS tissues were chopped into small pieces, incubated in RIPA for 2 h, and homogenized by pipetting every 20/25 minutes. Samples were then sonicated using Bioruptor Pico-Diagenode.

For cell cultures: Cells were harvested in PBS and centrifuged at 500 rcf for 3 minutes. PBS was removed by suction, and samples were resuspended in 80 μL of RIPA with protease inhibitor (Sigma-Aldrich Cat# P2714-1BTL). Cells were incubated for 15 minutes at 4 °C on an orbital shaker and then sonicated with Bioruptor Pico-Diagenode. After sonication, samples were centrifuged at 16,000 rcf for 5 minutes at 4 °C. Supernatant was kept, and BCA was used to measure the protein content of each sample.

### Western blot

Cell cultures and animal tissues: 15 μg of protein extracts were loaded on 10% pre-cast gels (Bio-Rad Cat.# 4568034) and run for 1hr at 100V. Gels were then transferred on PVDF membrane (Millipore Cat.# IPVH00010) overnight at 4 °C at 30V and developed with primary and secondary antibodies as indicated. Membranes were developed with SuperSignal™ West Femto Maximum Sensitivity Substrate (Thermo Fisher Scientific Cat# 34094), and imaged using ChemiDOC XRS+ System (Bio-Rad).

Human samples: The gels were U.V. activated for 5 minutes before transferring onto the 0.22 μm nitrocellulose membrane. The semi-dry transfer was done using a Trans-Blot Turbo Transfer System (Bio-Rad) at 25mV for 10 minutes. A stain-free blot image was captured after the transfer and used as total protein. The blot was processed as previously described. The quantification was done using Image Lab Software (Bio-Rad); the intensity of Kap β2-positive band is normalized to its total protein.

### Immunoprecipitation

Flag: Immunoprecipitation was carried out using anti-Flag agarose beads (mouse IgG agarose beads were used as control). Rat primary cortical neurons and tissues were extracted in IP buffer containing 50 mM Tris-HCl (pH 7.5), 150 mM NaCl, 1mM EDTA, 0.5% NP-40, 100 mM EDTA (pH 8.0) supplemented with protease inhibitor (Sigma-Aldrich Cat.# P2714-1BTL). After incubating at 4⁰C for 30 min., the samples were sonicated and then centrifuged for 10 min. at 10,000 rcf. Bradford assay was used to determine protein concentration. The agarose beads were added to 600 μg of cell lysates and incubated at 4 °C for 24h under constant shaking. Cell lysates were centrifuged for 1 min. at 1,000 rcf to collect the supernatant. The pellets were washed five times, but only the first wash was collected. After the last centrifugation, the samples were eluted with SDS-PAGE loading dye at 95 °C for 5 min. The entire eluate was run for Western blot. 2% of total lysate was used as input.

GFP and Kap β2: Immunoprecipitation was carried out using magnetic beads (Thermo Fisher Scientific Cat.# 88804) following manufacturer instructions. Briefly, 25uL of magnetic beads were conjugated with 10ug of the desired antibody overnight and then crosslinked for 30 minutes using DSS (disuccinimidyl suberate). The obtained beads-antibody complex was incubated with cell or tissue lysates overnight. The next day beads were washed and the bound protein was eluted. The entire eluate was run for Western blot. 2% of total lysate was used as input. For Kapβ2 IP double volume of beads was used.

### Immunofluorescence

Primary cortical neurons were fixed for 20 minutes at 37 °C with 4% PFA and incubated in PermBlock solution (2% BSA, 0.3% Triton-x, 5% donkey serum in PBS). Primary antibodies were diluted in 0.1% BSA cells and were incubated overnight at 4 °C on an orbital shaker. The following day cells were washed two times with PBS at room temperature for 10 minutes and incubated with the appropriate secondary antibody for 1h at room temperature. Cells were then washed twice in PBS and incubated with Hoechst solution (Thermo Fisher Scientific Cat.# 62249) for 10 minutes at room temperature to stain the nuclei. After a final rinse in PBS, coverslips were mounted on glass slides using AquaMount (Lerner Laboratories Cat.# 13800). Tissues were cryo-sectioned using Cryostar NX50. 20 μm thick sections were placed on glass slides and incubated for 10 minutes in 4% PFA. Slides were then washed in TBS-T for 10 minutes and incubated for 1h at room temperature in PermBlock solution (2% BSA, 0.3% Triton-x, 5% donkey serum in PBS) and then incubated with primary antibody diluted in 1% BSA, 0.3% Triton-x and 1% donkey serum overnight at 4 °C in a humidified chamber. The following day, slides were washed with TBS-T, incubated with the appropriate fluorescent secondary antibody for 1 hour at room temperature, rinsed twice in TBS-T, incubated for 10 minutes with Hoechst (Thermo Fisher Scientific Cat.# 62249), and mounted on coverslips using AquaMount (Lerner Laboratories Cat.# 13800) for imaging analysis.

### Antibodies and dilutions

Primary antibodies: anti-Kapβ2 (RRID:AB_2206884, Santa Cruz Biotechnology, Inc Cat.# sc-101539; WB dilution 1:1000); anti-Kap β2 (RRID:AB_262123, Sigma Cat.# T0825; IF dilution 1:500); anti-Map2 (RRID:AB_2138178 Novus Cat.# NB300-213; IF dilution 1:2000); anti-GFP (RRID:AB_11042881, Proteintech Cat# 50430-2-AP, WB dilution 1:1000); anti-TDP-43 (RRID:AB_615042, Proteintech Cat.# 10782-2-AP; IF dilution 1:500); anti-GAPDH (RRID:AB_2107436, Proteintech Cat.# 60004-1-Ig; WB dilution 1:1000); Neun (D4G40) XP(RRID:AB_2651140, Cell Signaling Technology Cat.# 24307; IF dilution 1:500): FUS (RRID:AB_2247082, Proteintech Cat.# 11570-1-AP; IF:1:500) Secondary HRP-conjugated antibodies: HRP-Conjugated anti-rabbit (RRID:AB_772191, Cytiva Cat.# NA9340-1ML; WB dilution 1:10,000); HRP-Conjugated anti-mouse (RRID:AB_772193, Cytiva Cat.# NA9310-1ML; WB dilution 1:10,000); HRP-Conjugated anti-rat (RRID:AB_2936877, Sigma Cat.# AP136P; WB dilution 1:3,000); Secondary fluorescent antibodies: AlexaFluor-546 anti-mouse (RRID:AB_2534012, Thermo Fisher Scientific Cat.# A10036; IF dilution 1:1,000); AlexaFluor-647 anti-rabbit (RRID:AB_2536183, Thermo Fisher Scientific Cat.# A31573; IF dilution 1:1,000); AlexaFluor-647 anti-chicken (RRID:AB_2762845, Thermo Fisher Scientific Cat.# A32933; IF dilution 1:1,000);

### Extraction of total RNA and quantitative Real-Time PCR

Total RNA was extracted using Trizol Reagent (Ambion Cat.# 15596026) following the manufacturer’s instructions. Briefly, cells were lysed in Trizol Reagent and homogenized using a syringe for insulin. RNA was then extracted in isopropanol. Samples were centrifuged for 15 min. at 12,000 rcf. The RNA-containing pellet was washed with 75% ethanol and centrifuged for 5 min. at 7,500 rcf. Dried pellets were resuspended in DEPC water, and concentration was assessed with Nanodrop. 500ng of total RNA was first treated with DNAse I (Thermo Fisher Scientific Cat.# EN0521) and retrotranscribed using SuperScript™ IV Reverse Transcriptase (Thermo Fisher Scientific Cat.# 18090010). The cDNA was diluted 1:2 and used for qRT-PCR, which was performed using the following TaqMan assay: GAPDH-VIC (Thermo Fisher Scientific Cat# 4352338E) and Kap β2-FAM (Thermo Fisher Scientific Cat.# 4331182 Assay# Rn01489969_m1). 50ng of cDNA was used per reaction. Samples were analyzed using Design and Analysis tool by Thermo Fisher Scientific. Results are shown as fold change using the 2 ^ΔΔCt^ method.

### Confocal Imaging Analysis

Confocal images were taken on a Nikon A1R confocal microscope with a 60X objective. Z-stacks were taken for each image. 0.2 μm Z-stacks were taken to perform colocalization studies and puncta counting using a built-in function in Imaris. One μm stacks were taken for TDP-43 nuclear intensity studies. Maximum intensity projection was then generated through a built-in function of the A1R analysis software. The nuclei of each co-transfected cell were selected as the region of interest, and TDP-43 intensity was assessed. To analyze aggregates content, the mask was made on the mCherry channel and the TDP-43 fluorescence was subsequently assessed.

FRAP experiment was performed in cells that were green (GFP or GFP-Kap β2) and red (mCherry, GR _50_-mCherry, GR _100_-mCherry) fluorescent. Aggregates were selected as regions of interest and photobleached for 2 sec. with blue light with laser power at 25% power. Following photobleaching, images were taken every 2 seconds for 3 minutes and analyzed with Nikon A1R software. Normalization and curve fitting was performed using the online tool EasyFRAP (normalization was performed using double normalization and curve fitting was performed using double equation). Representation of fluorescence recovery intensity curves over time were obtained using Prism GraphPad 9.0.

### Time-lapse microscopy

Cells were imaged every two days after transfection. 20 × objective was used, and 25 fields/well were acquired. Only cells positives for both Kap β2 and mCherry were counted. Cells were scored dead when they disappeared from the field of view, or clear signs of neurites fragmentation appeared. Kaplan Meier curves were then constructed in Prism GraphPad 9.0.

### Visualization of TDP-43 Condensates

His_6_-SUMO1-TDP-43 (5 μM) was incubated with an equimolar concentration of (GR) _20_ for 1 hour at room temperature in PBS. 200 nM His _6_-SUMO1-TDP-43-GFP and 100 nM TAMRA-(GR)_20_ were added to visualize TDP-43 and (GR) _20_. The reaction mixture was incubated at room temperature for 1h, and samples were spotted onto a coverslip and imaged by DIC and fluorescent microscopy using a Leica DMi8 Inverted microscope. To inhibit condensation, 5 μM Kapβ2 was added at the beginning of the assay.

### Proteomics analysis

Publicly available data sets from Liu, F., Morderer, D., Wren, M.C. et al. (Liu et al., 2022) were downloaded from massive.ucsd.edu (MassIVE MSV000088581). Analysis and statistical comparison of protein abundance were performed using basic packages (ggplot, ggplot2) of R (version 4.0.5) and Rstudio (version 2022.07.2).

### Transcriptomics analysis

In this study, we performed a transcriptomic analysis using publicly available datasets to investigate Kapβ2 expression patterns in iPS derived cortical neurons and ALS patient cerebral tissue. For the AnswerALS data: The raw sequencing data were processed using quality control metrics and aligned to a reference genome using Kallisto. Differential gene expression analysis was performed using DESeq2. Kap β2 expression levels across samples were analyzed with ggplo2 and statistical analysis were carried out using GGPUBR. For the human tissue data raw counts were downloaded from https://github.com/NathanSkene/ALS_Human_EWCE. Kapβ2 expression levels across samples were analyzed with ggplo2; statistical analysis were carried out using GGPUBR.

### Statistical analysis

All experiments were performed at least 3 times (n ≥3). At least 3 mice/group were used. For live-cell imaging, >150 neurons/condition were counted for each n. For colocalization, puncta counting, and TDP-43 intensity, ≥10 neurons/condition were counted for each n. For the FRAP experiment, ≥5 cells/conditions were analyzed for each n. When two groups were compared, the t-student test was performed using the built-in function in Prism GraphPad 9.0. The Kaplan Meier curves were compared by the log-rank and Cox proportional hazards tests, and the statistical significance between curves was assessed with Cox (GraphPad software). For the FRAP experiment, statistical differences between the curves were evaluated by curve analysis (one-way ANOVA test) in Prism GraphPad 9.0.

## FIGURE LEGENDS

**Supplementary Figure 1**

A. qPCR analysis of GFP levels in cortical neurons transduced with GFP or GFP-GR _50_. Data are represented as mean ±S.E.M. (n=6, Student t-test, n.s.)

B. qPCR analysis of GFP levels in cortex and spinal cord of GFP and GFP-GR_50_ mice. Data are represented as mean ±S.E.M. (n=3, One Way-ANOVA, n.s.)

C. Box plot showing Kapβ2 transcript levels in iCN derived from control (healthy) and C9-ALS individuals. (n>15, Student t-test, n.s.)

D. Box plot showing Kap β2 transcript levels in frontal cortex of control and C9-ALS patients (n>10, Student t-test, n.s.).

**Supplementary Figure 2**

A. Upper inset: confocal imaging of WT mouse spinal cord. Red is Kap β2; far red is NeuN. Lower inset Imaris rendering of Kapβ2 staining in WT mouse spinal cord.

**Supplementary Figure 3**

A. Venn diagram showing the protein interacting with GFP and GR_50_-GFP

B. Floating bar graph showing quantification of the intensities of the peptides for Kap β2 and Kapβ2b in GFP and GR_50_-GFP.

C. Western blot of the immunoprecipitation assay performed on GR _50_-GFP transduced cortical neurons. The same input was process with Flag-Beads, GFP-beads and Kap β2 beads. Blot were probed for GFP and Kapβ2. Total protein was used as loading control

D. Western blot of the immunoprecipitation assay performed on cortical neurons transduced with GR _50_-GFP virus. The input was process with Kap β2 beads. Blot were probed for GFP and Kapβ2. Total protein was used as loading control

E. Western blot of the immunoprecipitation assay performed on cortex of GR _50_-GFP mice. The same input was process with Kap β2 beads. Blot were probed for GFP and Kap β2. Total protein was used as loading control

F. Western blot of the immunoprecipitation assay performed on cortex of GR _50_-GFP mice with anti-goat and anti-rat Igg control. Upper panel: The same input was process with Kap β2 beads and anti-rat Igg. Blot were probed for Kap β2. Lower panel: The input was process with anti-goat Igg. Blot were probed for GFP.

**Supplementary Figure 4**

A. Confocal images of primary rat cortical neurons treated with Kap β2 siRNA 100 μM or scramble. FUS is in green, Map2 is in magenta and DAPI is in blue. Scale Bar is 5μm.

B. Bar graph showing the quantification of nuclear intensity of FUS Data are represented as mean ±S.E.M. (n=3, Student t-test, n.s.)

**Supplementary Figure 5**

A. Representative images of censored neurons over time transfected with 200/400ng of Tm^+^/GFP^+^. Green is Kapβ2; red is Td-Tomato. Scale bar 20μm.

B. Kaplan-Meier curves of time-lapse experiments on rat primary cortical neurons transfected with 200/800ng of Tm ^+^/GFP-Kapβ2^+^ (GFP used as control) per 150,000 cells. Neurons double positive Tm ^+^/GFP^+^ were counted (n=3, m>150 neurons, Log-rank Mantel-Cox test: n.s.)

**Supplementary Figure 6**

G. Epifluorescence images of condensates formation in the presence of TDP-43, (GR)_20,_ and BSA. Green is TDP-43, and red is GR_20_. Scale bar: 15μm.

