## Supplementary figures and images for "The nuclear import receptor Kapβ2 modifies neurotoxicity mediated by poly(GR) in C9orf72-linked ALS/FTD"

### Supplemental Figure 1

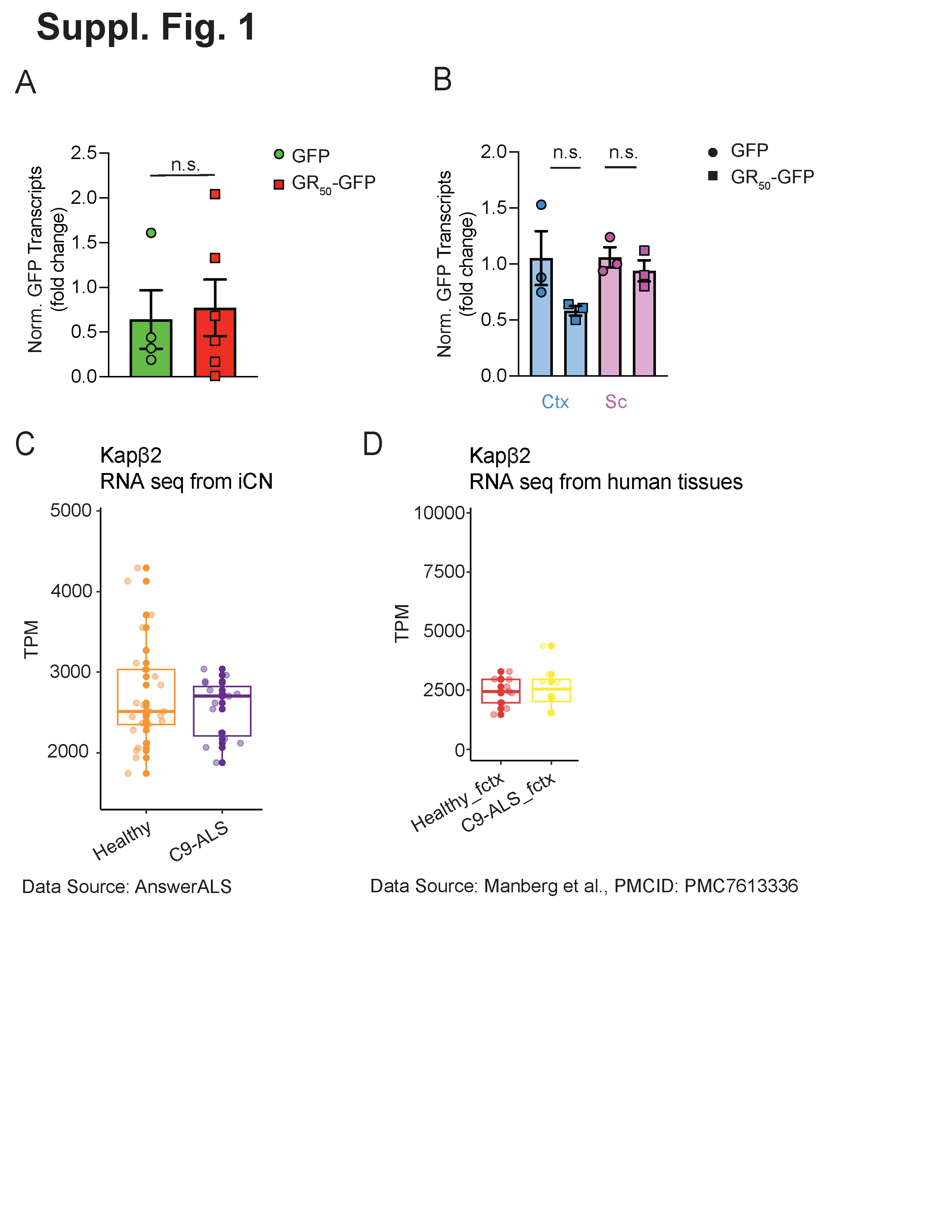

### Supplemental Figure 2

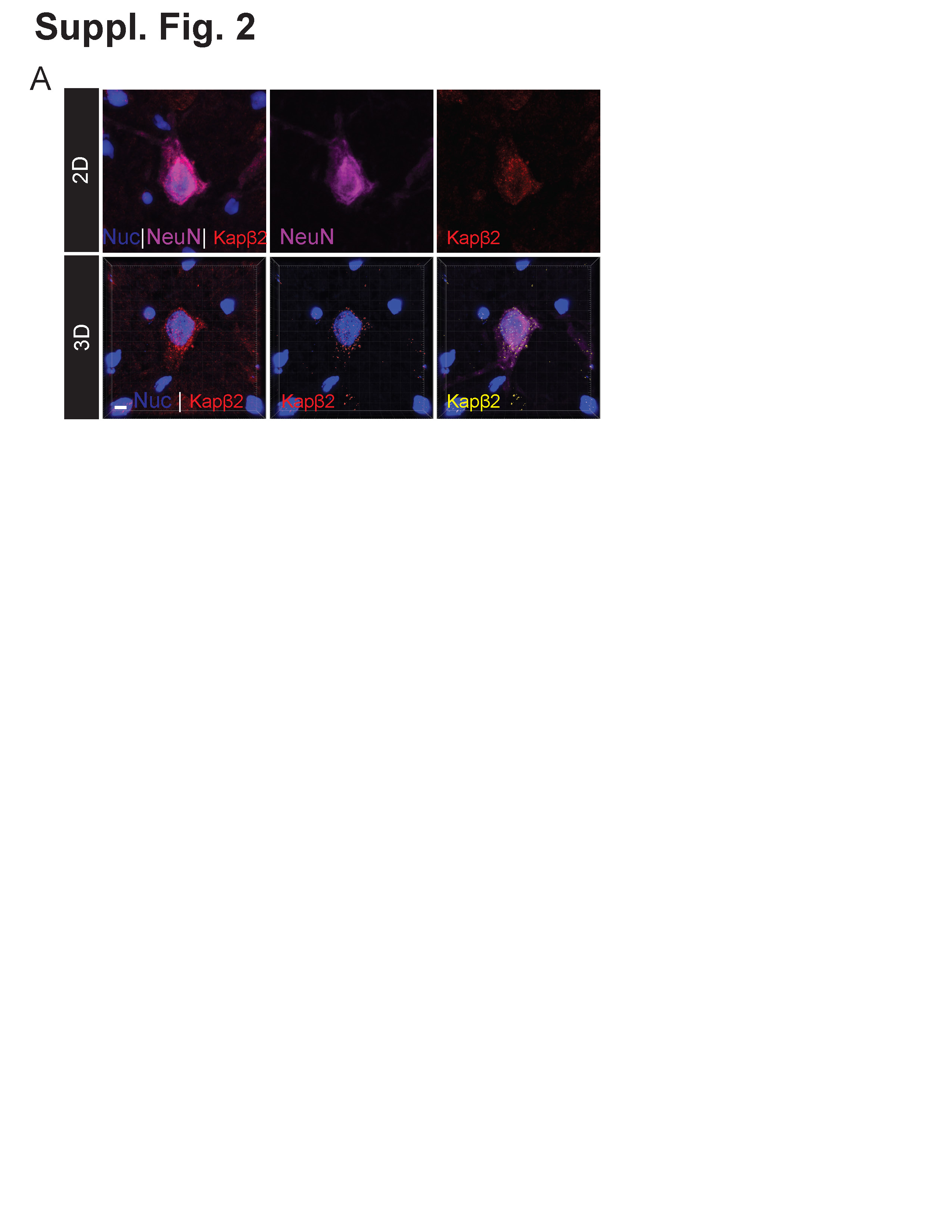

### Supplemental Figure 3

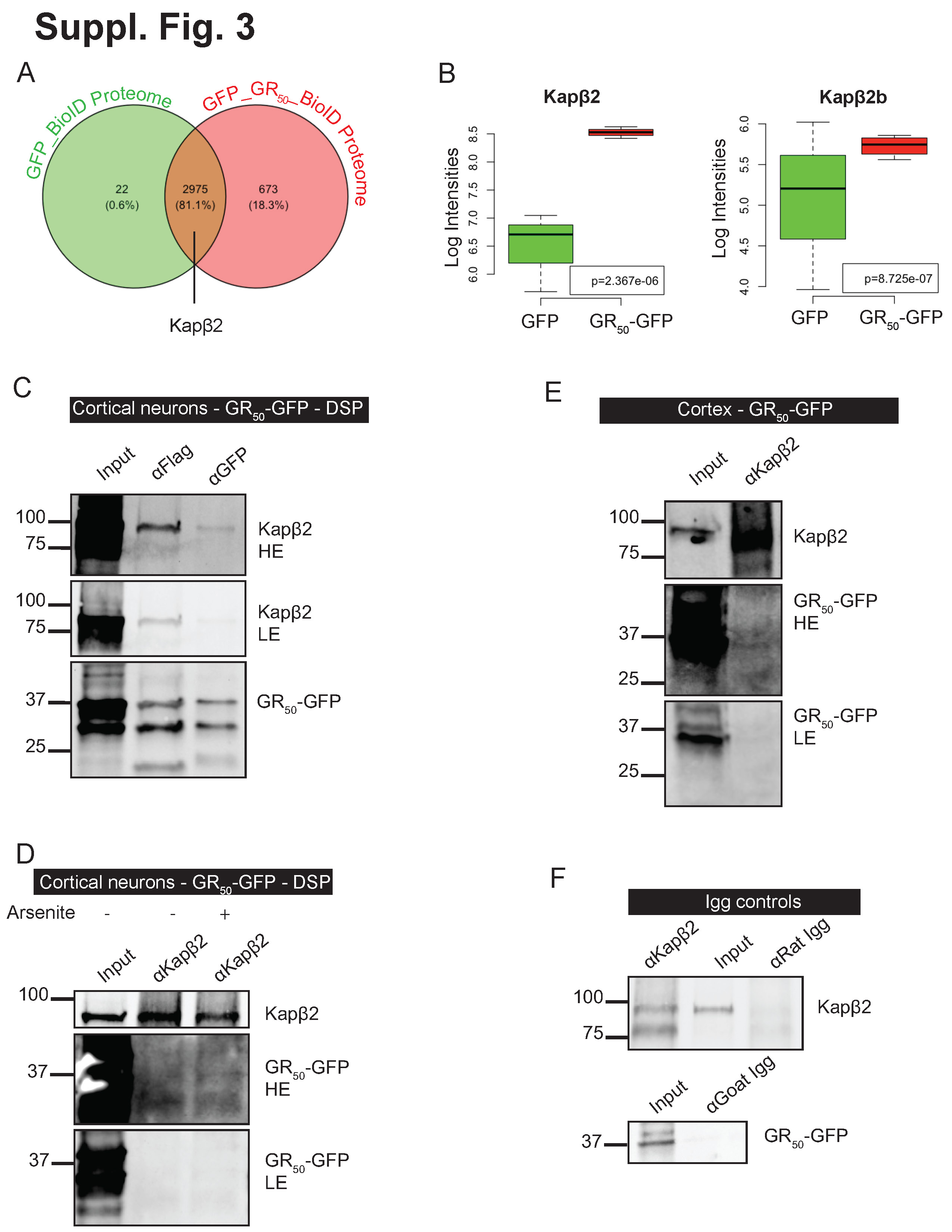

### Supplementary Figure 4

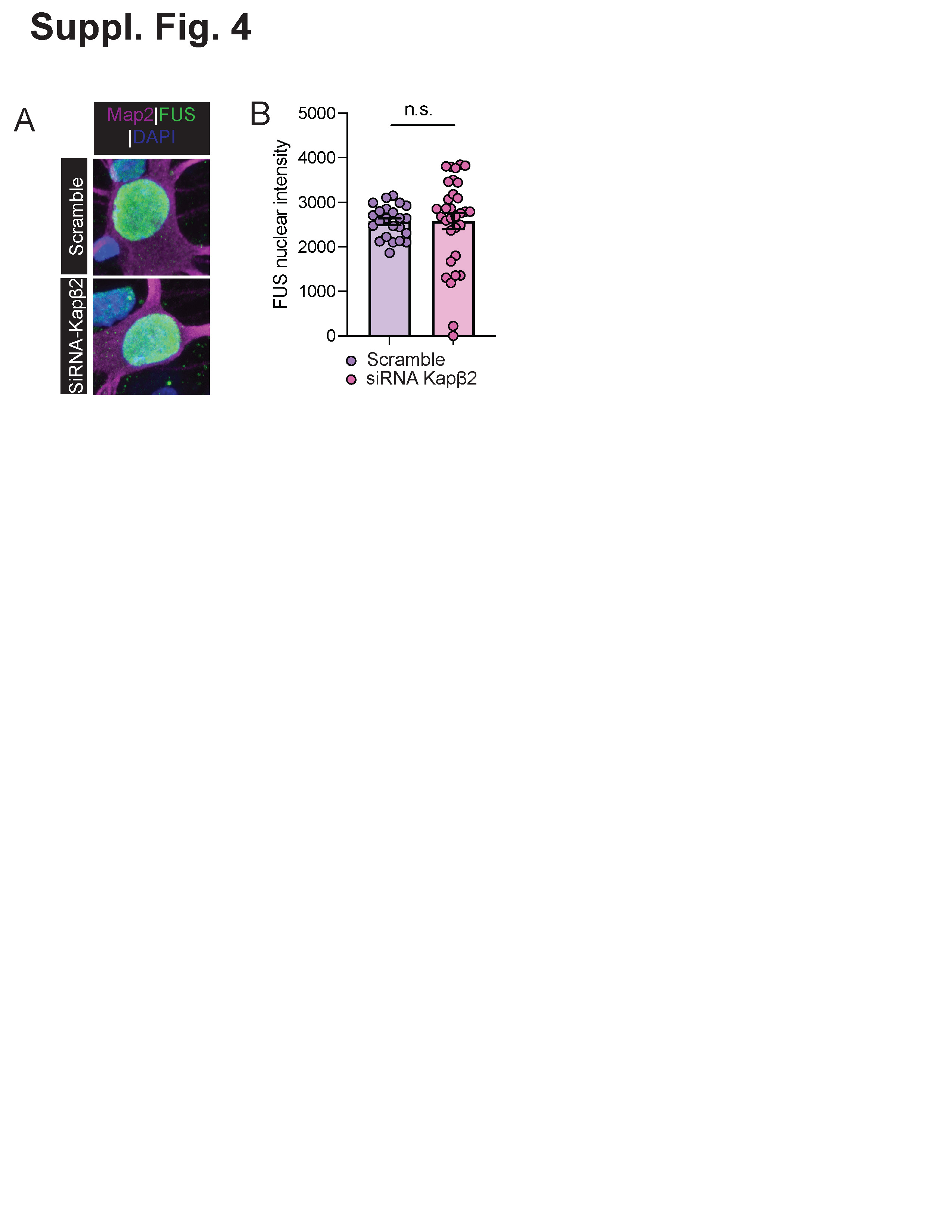

### Supplementary Figure 5

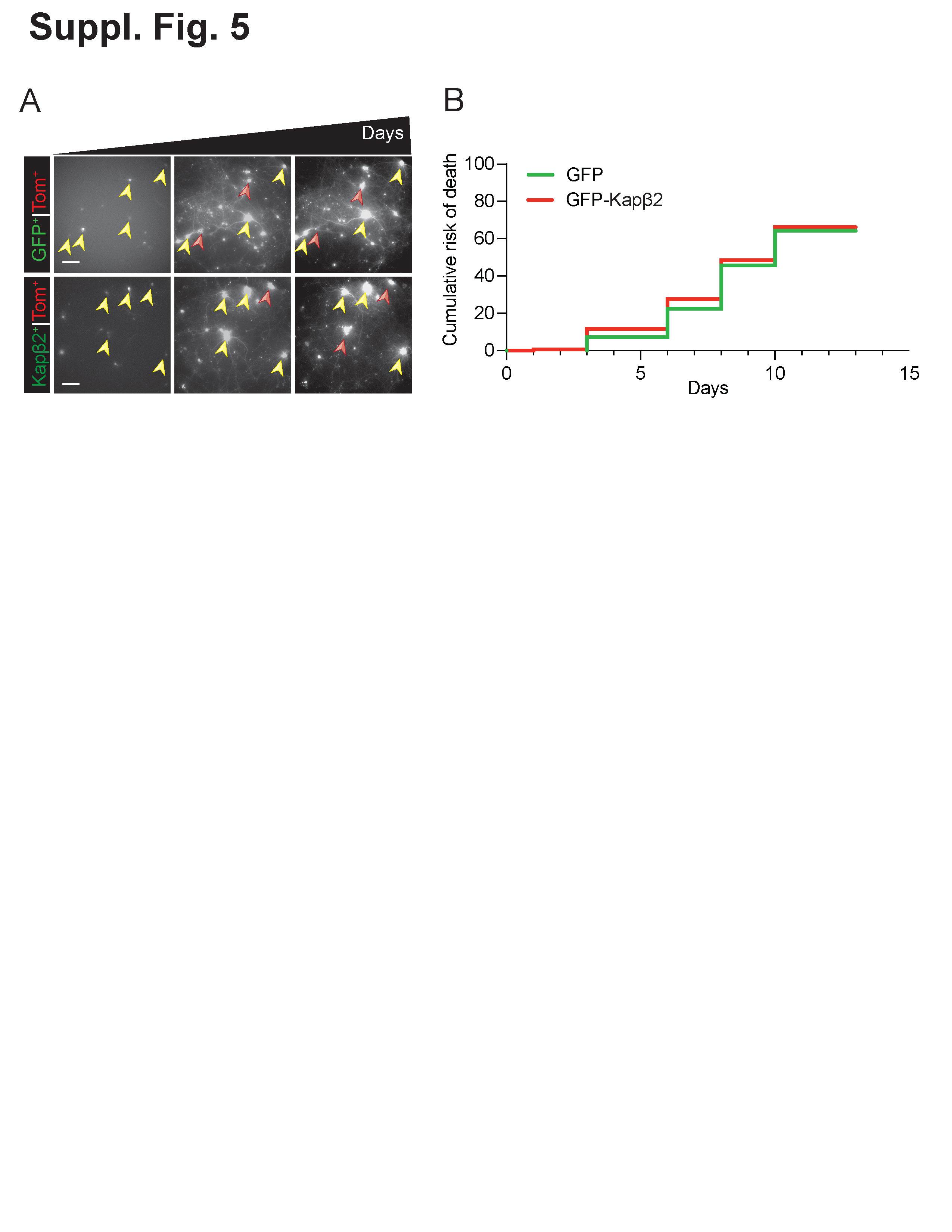

### Supplementary Figure 6

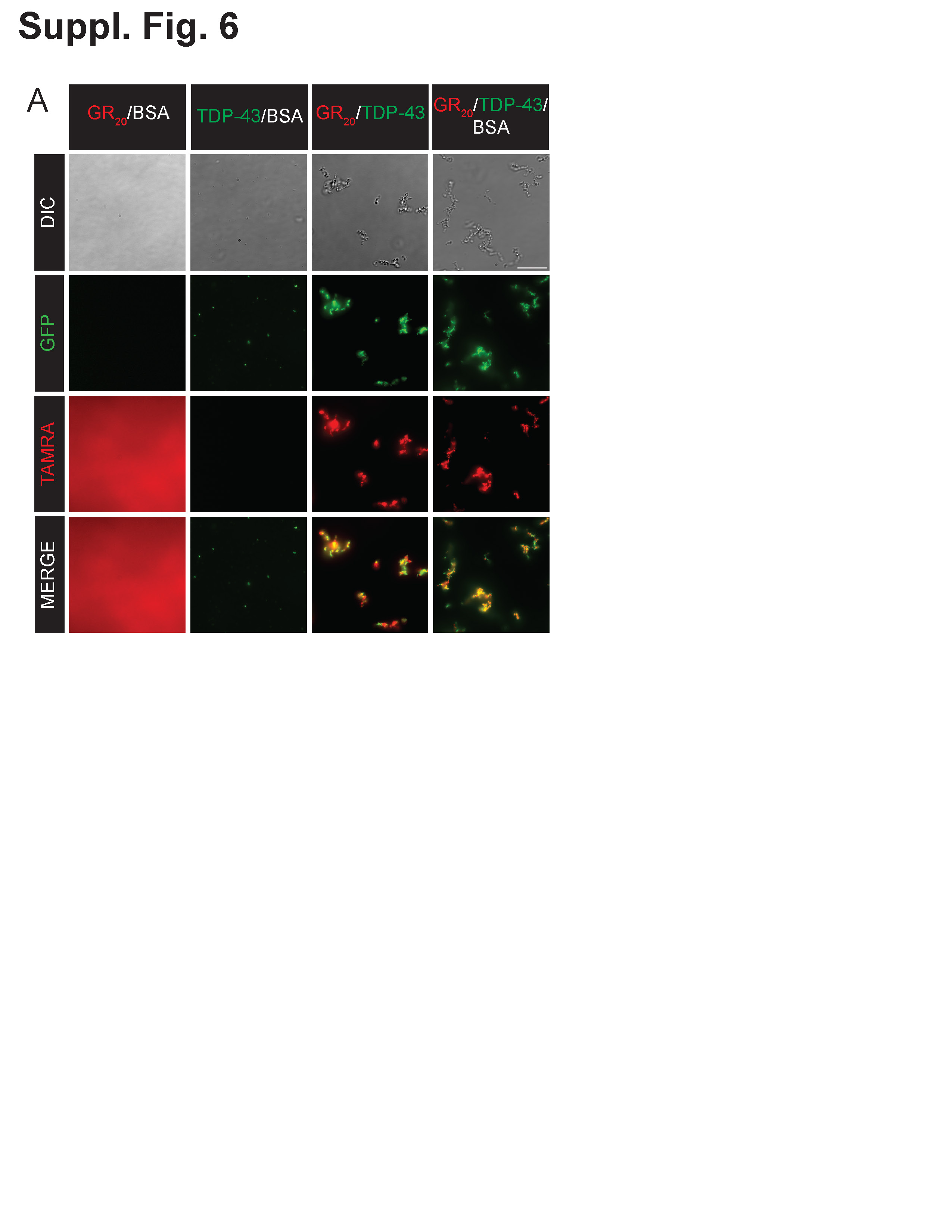
